# Cell-Free Recombinase Integrated Boolean Output System (CRIBOS)

**DOI:** 10.1101/2024.10.22.619686

**Authors:** Jingyao Chen, Aviva Borison, Douglas Densmore, Wilson W. Wong

## Abstract

Cell-free gene expression systems are increasingly important in fundamental research and biomanufacturing, offering a versatile platform for studying gene circuits and bio-computation. We present the <u>C</u>ell-Free <u>R</u>ecombinase Integrated <u>B</u>oolean <u>O</u>utput <u>S</u>ystem (CRIBOS), a site-specific recombinase-based multiplex genetic circuit platform designed for cell-free environments. With CRIBOS, we build over 20 multi-input-multi-output circuits, including 2-input-1-output genetic circuits and a 2-input-4-output decoder. Combined with allosteric transcription factors (aTFs)-based sensors, the circuits demonstrate multiplex environmental sensing. Moreover, utilizing paper-based CRIBOS, which demonstrates remarkable portability and stability in a paper-based setting, we present a biological memory storage device that can preserve DNA-based biological information for over four months with minimal resources, energy costs, and maintenance requirements. Implementing CRIBOS not only expands the application of multiplex Boolean logic gates from cellular systems to the cell-free environment but also augments their overall versatility, opening new avenues for designing and applying sophisticated genetic circuits.

## Introduction

Genetic circuits are foundational tools in synthetic biology with applications in medicine^1,2^, diagnostics^3–5^, agricultural production^6,7^, and environmental surveillance^8,9^. Synthetic gene circuits enable dynamic, precise, interactive, and logical control of cell function, which can facilitate drug discovery by enabling real-time monitoring of diverse molecular events and disease-relevant dynamics^1^. Genetic circuits have also been applied to develop new classes of rapid, cost-effective, deployable diagnostic devices applicable to various diseases and chemicals^4^. Moreover, in chemical manufacturing, engineered cell factories equipped with sense-and-response systems enable dynamic control of expression pathways and optimal allocation of cellular resources^10^. In agriculture, crop yield can be boosted by smart plants incorporated with synthetic circuits capable of sensing and adapting to environmental challenges^11^.

### Recently, cell-free systems have been increasingly recognized as valuable platforms for fundamental interrogation and biomanufacturing^12–14^

A cell-free system is a membrane-free, simplified version of cells that operates without concern for cell viability. It provides many advantages for developing and characterizing circuits and bio-computation, such as a large dynamic range for chemical measurements^15^, reduced interference from irrelevant cellular pathways^16^, and a high level of control over the composition of the genetic circuits^16,17^. Moreover, a cell-free environment can detect small molecules that are toxic or impermeable to cells^18,19^. It can provide clearer signals and exclude most of the intrinsic noise caused by complex organelles and biochemical reactions inside the cell by simplifying the protein composition^20^.

With its portability and ease of detection, cell-free systems have been combined with diverse genetic circuits for point-of-care diagnostics^21^, environmental sensing^22^, and fundamental research on metabolic pathways^23,24^. Examples include but are not limited to the aTFs-based water contaminant detection system ROSALIND (RNA Output Sensors Activated by Ligand Induction)^22^, as well as the CRISPR-based infectious disease diagnostics device SHERLOCK and DETECTR^25^. These examples demonstrate the high sensitivity and robustness of genetic circuits created in cell-free systems and provide an ideal platform for research in synthetic biology and the construction of bio-computing devices.

Boolean algebra, with its long history in computing, offers a powerful framework for bio-computing due to its expressiveness and the efficiency of its well-understood algorithms^26^. Boolean algebra has been foundational in the design and analysis of digital circuits, forming the basis of computer logic and programming^26,27^. Boolean logic gates, such as AND, OR, and NOT, can describe intricate computational tasks and can be transformed into equivalent functionalities efficiently. This makes it particularly suitable for constructing complex genetic circuits in synthetic biology, where precise control and predictability are paramount^28,29^. Various types of bio-computing devices and Boolean logic gates have been applied to different bacterial^30^, yeast^31^, mammalian cell strain^32,33^, and animal models^34^ to enable complex bio-computation. However, their implementation in the cell-free field is not as developed, and circuits with multiple inputs and multiple outputs are still limited in the cell-free system, primarily due to limited tools for building cell-free biocomputing circuits. This limitation necessitates the integration of multiple layers of genetic logic gates to construct biocomputing devices and requires extensive construction and characterization for circuit design. The restricted exploration and construction of cell-free Boolean logic gate toolkits hinder the broader application of cell-free systems. This paper creates a cell-free Boolean biocomputing platform for complex multi-input-multi-output genetic circuit constructions. It makes a streamlined procedure to simplify the construction of cell-free genetic circuits to facilitate the study of complex metabolic pathways, controlled biologics manufacturing, and multiplex environmental sensing.

### Site-specific DNA recombinases are powerful tools for developing multiplex Boolean logic gates in various organisms and models

such as bacteria^35^, mammalian cells^36,37^, and mouse brains^38^. In addition, they perform different types of modification, such as excision and inversion, based on the direction of the recognition sites. Moreover, many recombinases and their recognition sites have been developed or discovered, each targeting their corresponding recognition sites with high efficiency and orthogonality, further expanding the pool of usable genetic tools^36^. The success of recombinase-based genetic circuits in various living organisms proves that recombinases can be easily adapted to different platforms. However, recombinase-based Boolean logic gates have yet to be applied to a cell-free system.

### To expand the toolkit for genetic circuits, we developed a cell-free recombinase system

We demonstrated the ability to engineer complex circuits that can be easily implemented and function without trial and error. Although recombinase has been widely studied and well-characterized in a cellular system, transitioning from cellular hosts to cell-free systems is not simple. To apply the recombinase circuits to cell-free systems, we comprehensively characterized circuit designs and established design rules applicable to cell-free genetic circuit development.

Additionally, the irreversibility of recombinase excision can produce a DNA writer for biological memory recording. With its high fidelity, recombinase has been previously applied to cellular systems to develop DNA-based memory devices, such as BLADE and Recombinase State Machine (RSM)^35,36^. The successful, sophisticated tools illustrate that recombinase is an ideal candidate for implementing genetic circuits in a DNA-based memory storage device. However, while recombinase circuits have been used to implement DNA memory, they have yet to be explored in a cell-free environment. Thus, we also establish the potential of cell-free recombinase to develop DNA-based memory storage and characterize the stability of memory and biological information stored in such circuits.

## Results

### Design Characterization of Recombinases in a Cell-free Environment

Although recombinase genetic circuits have been widely applied to different cellular and animal models, exploration of their application in cell-free conditions is rare. As such, we have limited knowledge of their design rules. To address this issue, we characterized Cre recombinase excision circuits to establish the potential of recombinase in a cell-free reaction. We chose to use the Protein Synthesis Using Recombinant Elements (PURE) system because of its high purity, simple composition, easy manipulation, and robust performance in building genetic circuits for diagnostics and environmental surveillance.

The excision reporter comprises a T7 promoter, a ribosome binding site (RBS), a terminator flanked by two Cre recognition loxP sites, and a mCherry reporter gene (**Fig. 2A**). For the starting reporter design, we used a combination of rrnB T1 and T7Te terminator (Term 1), a double terminator combination verified in a bacterial recombinase system in the previous study, to suppress reporter expression before recombination^39^. In addition, the expected reporter plasmid post-excision, referred to as a LoxP-mCherry plasmid, was also generated. This serves as a positive control for the successful expression of the reporter plasmid after excision by Cre (**Fig. 2A**). With the expression and activity of Cre recombinase, we expect the expression of the mCherry reporter gene to go from OFF to ON, while the LoxP-mCherry should constitutively produce high reporter output. However, opposite results were observed. The reporter-alone condition showed a basal output signal that is a 27-fold change over the blank condition. In addition, an 82% and a 55% decrease in translational output were observed for reporter and LoxP-mCherry, correspondingly, when incubated with Cre-expressing plasmids. (**Supplementary** Fig. 1A). These results suggest multiple issues of the current circuit design. First, the high reporter basal expression indicates that the terminator between loxP sites was too weak and failed to terminate the T7 RNA polymerase activity. Second, the decreased reporter expression in the reporter+Cre condition indicates that Cre protein expression negatively impacts reporter expression, with the underlying mechanism remaining unclear.

**Figure 1.**
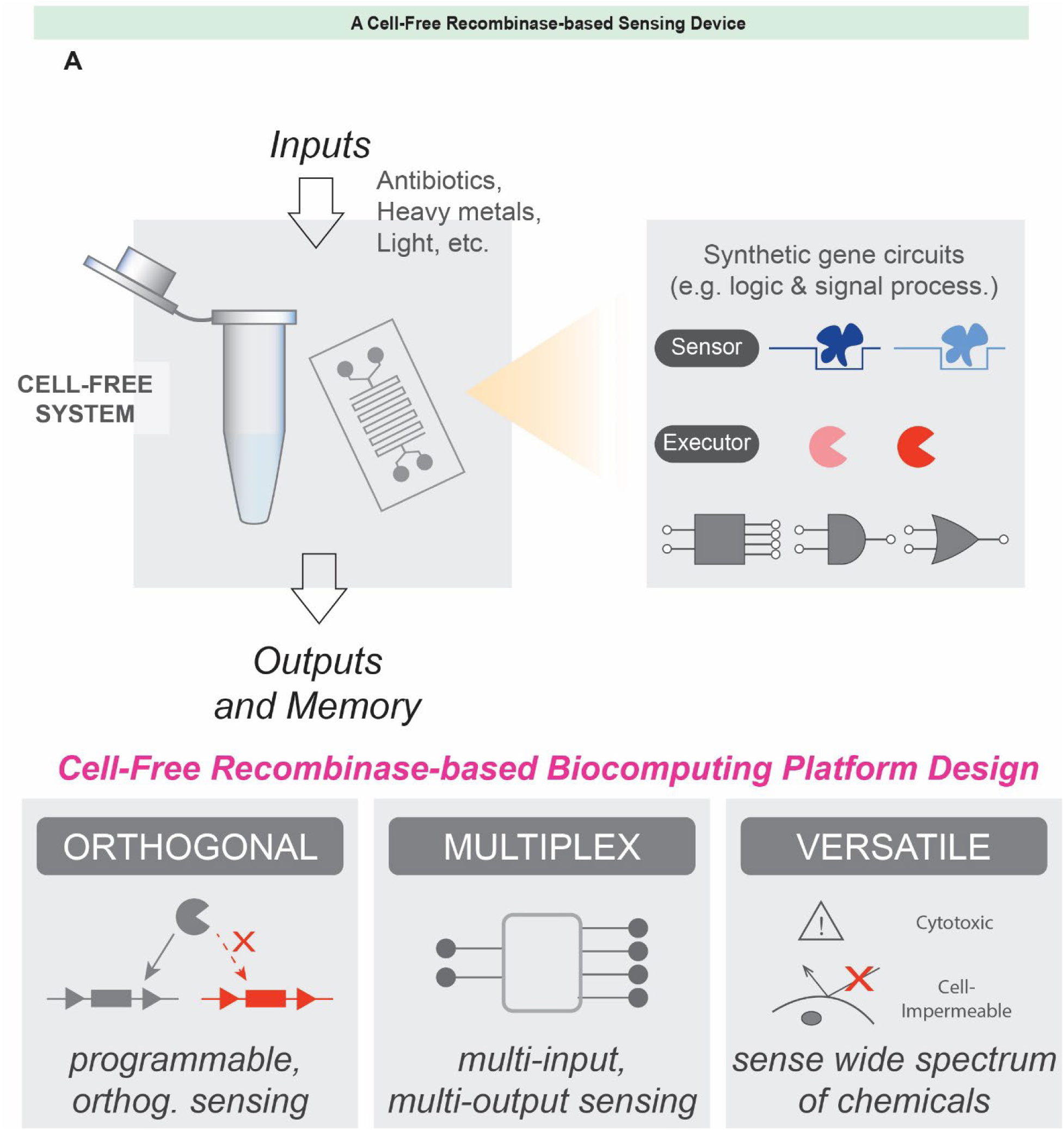
A cell-free recombinase-based biocomputing platform in test tube/microfluidics allows for the construction of orthogonal, multiplex, and versatile devices. Such a platform has potential applications in multiplex environmental sensing, biological information storage, and disease diagnostics.

**Figure 2:**
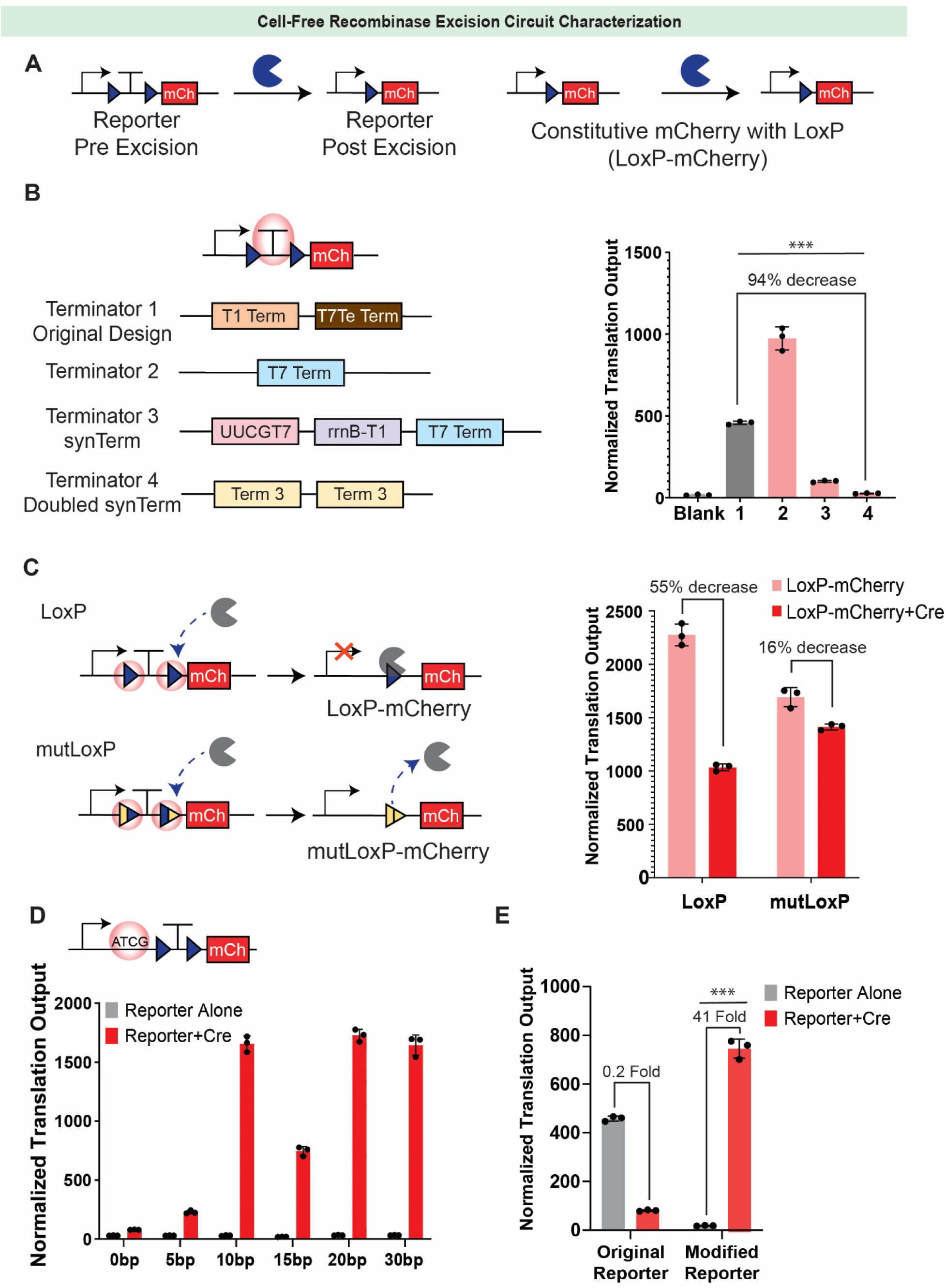
Characterization of cell-free recombinase excision circuits. (A) Excision reporter and LoxP-mCherry plasmids design. Before Cre excision, the terminator after the promoter suppresses transcription of the mCherry reporter gene. When Cre is expressed, the recombinase will excise the terminator and allow for transcription, thus triggering a mCherry fluorescent output. LoxP-mCherry plasmid, an expected reporter sequence post-excision, is cloned as a positive control for excision circuit testing. Blue triangle: loxP recognition site for Cre recombinase. T: transcription terminator. (B) Terminator optimization to prevent leaky reporter expression (left). mCherry expression of the pre-excision reporter (right). (C) Comparison of LoxP and mutLoxP sites on the reporter (left) and the corresponding mCherry output (right). (D) Normalized transcription output (NTO) as a function of insert length between promoter and recognition sites. (E) NTO comparison of original excision reporter and modified reporter. Data shown are the means of technical triplicate samples with error bars indicating +1 standard deviation. P-values were calculated as described in the methods.

We first focused on reducing the basal mCherry expression by utilizing strong terminators to limit the T7RNAP readthrough. We chose the T7 terminator, an engineered terminator for the T7RNAP expected to prevent T7RNAP readthrough efficiently, as the first candidate (Term 2). However, the results show that the T7 terminator also failed to thoroughly terminate the reporter transcription pre-excision (**Fig. 2B**). Our result agrees with a previous study of synthetic terminator design, in which the T7 terminator only showed 80% termination efficiency^40^. In addition, the previous research also screened an array of terminator designs and created a synthetic terminator (Term 3 - synTerm) that can prevent 99% T7RNAP readthrough. The synTerm is composed of a synthetic T7 terminator, rrnBT1 terminator, and T7 terminator (**Fig. 2B**)^40^. Replacing the terminator in our original reporter design with the synTerm reduced reporter leaky expression by >78%. Adding one extra copy of synTerm in the reporter (Term 4 - 2XsynTerm) can further lower 94% of reporter basal expression to a level similar to the blank condition (**Fig. 2B**).

With the basal expression issue solved, we next investigated the mechanism of the decreased reporter expression in the reporter+Cre condition. We hypothesized that the Cre binds to the loxP on the reporter post-recombination, thus blocking the T7 transcription. One loxP site will be left in the reporter plasmid after excision, thus serving as a repressive operator site for Cre to inhibit the promoter activity. To decrease the Cre binding to loxP sites post-excision, the loxP sites were replaced by the asymmetric, mutant loxP (mutLoxP) sites lox66/lox72. The lox66/72 pairs carry 5bp of mutation on the left or right end of the lox66 or lox72 sites, respectively, resulting in 10bp mutation on the recombined loxP sites post-excision (**Fig. 2C**)^41^. We hypothesize that the 5bp sequence mutation isn’t strong enough to prevent Cre recombinase from binding and modifying the DNA but will result in a 10-bp mutation post-excision, which can effectively prompt Cre recombinase to dissociate from the reporter plasmid. To test this hypothesis, we created a mutLoxP-mCherry plasmid by replacing the original loxP site with the mutLoxP site. Both LoxP-mCherry and mutLoxP-mCherry plasmids were then exposed to Cre recombinase to assess the impact of Cre expression on constitutive mCherry expression. Implementing the mutated lox66/72 sites recovers over 39% of reporter expression in the LoxP-mCherry plasmids (**Fig. 2C**).

The success of recovering output signal with the mutLoxP site suggests potential interference between T7 RNA polymerase and Cre binding to DNA. Recognizing the possibility of competition between these enzymes for binding, we hypothesized that the insert length, or distance between the promoter and the loxP sites, could impact the excision circuit expression. Thus, we also varied insert lengths to elucidate their impact on circuit performance. Among all the insert lengths that were tested, 10bp provides the most significant enhancement of reporter expression with and without Cre plasmid (**Fig. 2D**). We rationalize that it might be due to the competitive binding between the Cre recombinase and the T7RNAP – the two enzymes might interfere the other’s binding efficiency and activity if their binding sites are too close. However, ribosome binding sites should not be too far downstream to the T7 promoter since a longer sequence behind the transcription starting site will result in a longer non-coding mRNA sequence with a more complex secondary structure that can negatively affect translation efficiency. Thus, finding an optimal distance between promoter and recognition sites is critical, and 10bp is determined to be the insert length that can maximize the translation efficiency and T7RNAP activity in our system.

A cell-free system is supplemented with limited energy and enzymes. Thus, to better leverage the limited sources in the genetic circuits, we fine-tuned the ratio of reporter and recombinase plasmids for the excision circuits (**Supplementary** Fig. 1C). The dose-response curve of recombinase plasmids demonstrates that while a sufficient amount of recombinase plasmids is needed to trigger reporter signals efficiently, high-level of recombinase plasmids will also lead to decrease in reporter fluorescence. We hypothesize that this is due to the rapid depletion of energy and resources for recombinase expression, leaving insufficient resources for reporter fluorescence expression post recombinase excision.

With the characterization of the terminator sequence, recognition sites, insert length, and plasmid dosage, we present a cell-free Cre excision circuit that starts with a 0.2-fold induction, eventually optimized to 41-fold induction, resulting in more than 200X improvement of the excision reporter output signal (**Fig. 2E and supplementary Fig. 2**). This systematic optimization approach was also applied to excision circuits of five other recombinases in the cell-free environment. These recombinases have been proven to function in bacterial systems and can trigger above 20-fold change of reporter output signal with high efficiency and orthogonality in cell-free systems (**Supplementary** Fig. 1B).

### Environmental inputs for cell-free recombinase genetic circuits

Next, we interfaced the cell-free recombinase genetic circuits with biochemically relevant inputs using allosteric transcription factors (aTFs) to regulate T7RNAP activity. To control the transcription of recombinase, the cognate operator sequence is placed after the promoter. We reason that the binding of aTFs to the operator will prevent T7RNAP from accessing the promoter. In contrast, ligand binding will result in a conformation change of the aTFs, causing its release from the operator and thus triggering transcription of recombinase and the downstream excision circuit (**Fig. 3A**).

**Figure 3.**
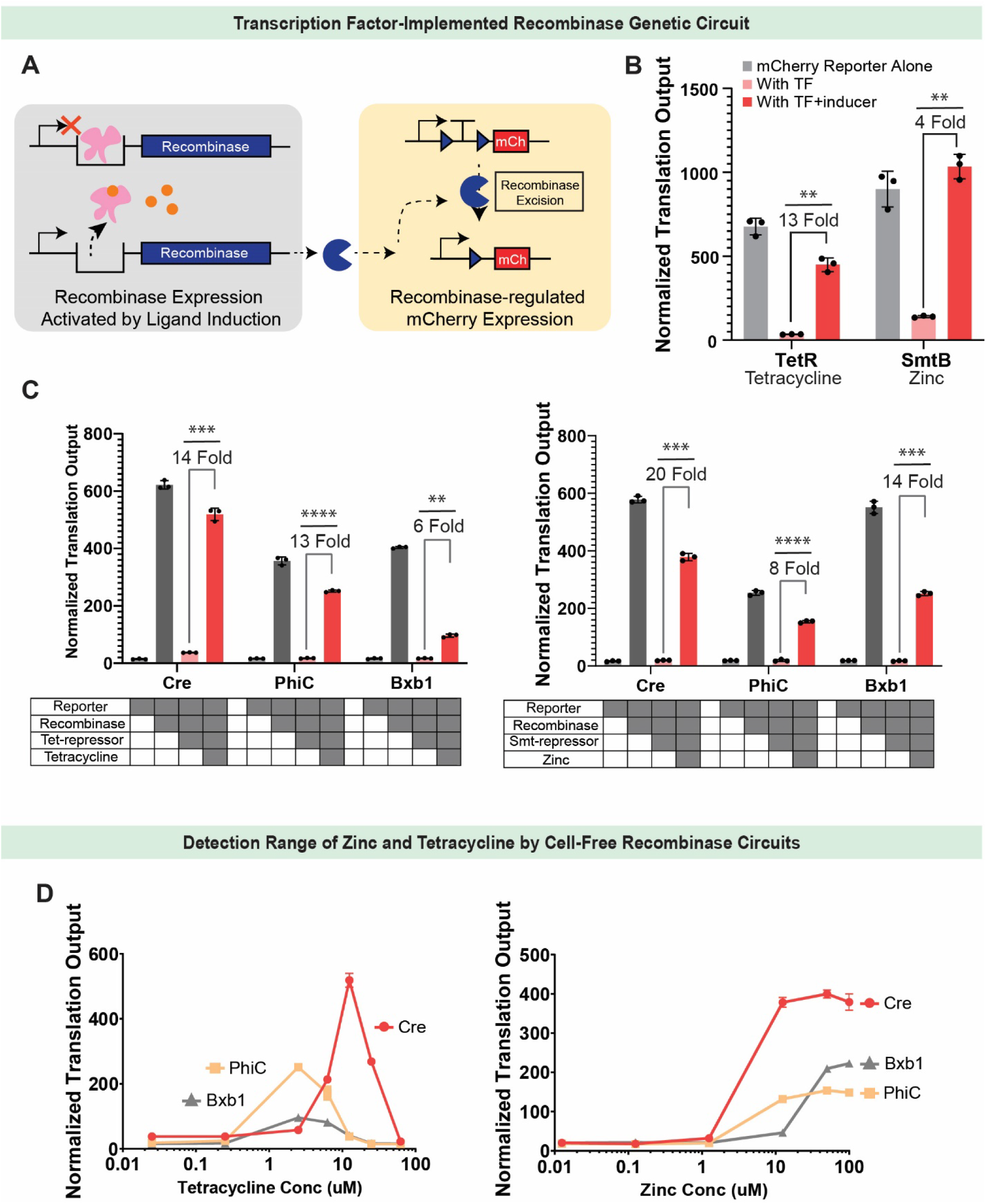
Characterization of aTFs-implemented recombinase genetic circuits in a cell-free condition. (A) Mechanism of aTFs-controlled recombinase genetic circuits. aTFs can bind to transcription operators behind T7RNAP and suppress transcription of the recombinase gene. Corresponding inducers bind to aTFs to release them from the operators, thus triggering recombinase expression and genetic circuit activation. (B) Characterization of tetR and smtB aTFs. (C) Cre, PhiC, and Bxb1 excision circuits controlled by tetR (left) and smtB (right) aTFs are tested. To set up the experiment, aTFs were produced, purified from bacteria, and added to the PURE reaction with genetic circuit plasmids with and without chemicals. (D) Chemical detection range is determined for each aTFs-implemented recombinase genetic circuit under optimal aTFs for each recombinase. To set up the experiment, 12.5uM tetR protein is used for the Cre excision circuit, 1.25uM tetR protein is used for PhiC and Bxb1 excision circuits, 10uM smtB protein is used for Cre, PhiC, and Bxb1 excision circuits. Data shown are the means of technical triplicate samples with error bars indicating +1 standard deviation. P-values were calculated as described in the methods.

We chose a group of aTFs previously tested and characterized in an *in vitro* transcription reaction, whose reaction environment is similar to that of a cell-free system^22^. To verify that the aTFs-operator interaction can efficiently control the expression of a mCherry reporter, an operator was inserted after the T7 promoter, following the design characterized in the previous study^42^. Within the seven pairs of aTFs-operator that verified to function in the cell-free environment, we selected well-studied tetR and smtB due to their highest yield and best performance (**Fig. 3B and Supplementary** Fig. 4A). Their corresponding cognate operators were then inserted into the Cre, PhiC, and Bxb1 plasmids to control recombinases expression (**Fig. 3C**). Dose-response curves for aTFs were performed to determine optimal aTF concentrations (**Supplementary** Fig. 3A).

After the optimal aTFs concentration was determined, a dose-response curve for small molecule inducers was performed to determine the detection window of the recombinase-based sensing circuits (**Fig. 3D**). It is essential to consider that the detection window for inducers varies among recombinase genetic circuits due to differences in their activities. For instance, stronger recombinases such as Cre necessitate higher concentrations of aTFs to fully deactivate circuit activity. As a result, these circuits exhibit a detection window that is shifted to the right compared to weaker recombinases like PhiC and Bxb1, detecting higher dosages of inducers. The observed shift in the detection window can be attributed to variations in aTFs concentration requirements, as different recombinase circuits demand distinct concentrations of aTFs. This diversity in recombinase activities allows us to broaden the detection window for inducers, as illustrated in **Figure 3D**. Furthermore, we assessed the limit of detection (LoD) for both zinc and tetracycline-induced sensors to determine the input dose that achieves detection above negative controls (**Supplementary** Fig. 3C). The fold change was calculated to determine the maximum change in reporter output. We found the limit of detection (LoD) for zinc-induced recombinase sensing was 1.17 uM for Cre, 2.69 uM for PhiC, and 4.45 uM for Bxb1, with fold changes of 20.7x, 8.14x, and 12.8x respectively. These values indicate that for zinc-induced sensing, Cre achieves the highest fold change in reporter response and detection at the lowest dose of zinc input. For tetracycline-induced sensing, the detection window was between 1.08 – 59.0 uM for Cre, 0.254 – 19.6 uM for PhiC, and 0.366 – 22.5 uM for Bxb1, with fold changes of 14.0x, 14.6x, and 5.76x respectively. These results show that while PhiC has an initial lower limit of detection, Cre has the broadest range in detectable tetracycline. Moreover, it was also observed that tetracycline at high dosages negatively affected the reporter gene expression (>12.5uM). This is most likely due to tetracycline’s inhibition of bacterial protein synthesis by preventing the association of aminoacyl-tRNA with the bacterial ribosome^43^. Additionally, we plotted the ROC curves and calculated AUC to quantify assay performance across sensitivity and specificity classification thresholds (**Supplementary** Fig. 3B). For zinc-induced sensing, we found AUC values of 0.907 for Cre, 0.833 for PhiC, and 1 for Bxb1. For tetracycline-induced sensing, the ROC curve analysis showed AUC values of 0.857 for Cre, 0.730 for PhiC, and 0.841 for Bxb1. These results demonstrate the promising ability of the assay to distinguish both true positive and true negative results.

### 2-Input-1-Output Logic Gates

The high orthogonality of recombinase and recognition sequences allows for more complex genetic circuits with multiple inputs and outputs to be developed under cell-free conditions, such as the 2-input, 1-output Boolean logic gates. We created twelve 2-input Boolean logic gates by placing Cre and PhiC recombination sites, lox66/72, PhiC attB/attP, respectively, by either side of terminator sequences and a mCherry reporter gene. In the original experimental design, the reporters and recombinase plasmids were premixed and added to the cell-free reaction before incubation. With this design, some circuits do not behave as expected in the logic table, which is mainly due to the delayed shutoff of the reporter fluorescent expression by PhiC recombinase (**Supplementary** Fig. 5). We reason that this is because of the weak activity of the PhiC recombinase. While Cre is very efficient and active and thus can promptly shut the reporter expression, the PhiC is much weaker and takes longer to excise the reporter gene. During this time, the reporter expresses and accumulates in the reaction simultaneously, leading to a leaky output signal for the genetic circuits. Therefore, we hypothesize that rapid recombination by PhiC is the key to decreasing the leaky signals, and pre-expression of the recombinases might help to achieve this goal.

In the following experimental design, instead of adding recombinase and reporter plasmids simultaneously to the cell-free reaction, we first pre-incubated Cre and PhiC recombinase plasmids in the cell-free reaction for 30 minutes. Then, we added the reporter plasmids to finish the circuit setup. The 2-input-1-output logic gates yielded intended behaviors (**Fig 4B and Supplementary** Fig. 6). It demonstrates that timing is also a critical factor in developing and optimizing genetic circuits, and a cell-free platform provides higher control and flexibility of timing over every step of circuits’ execution.

**Figure 4.**
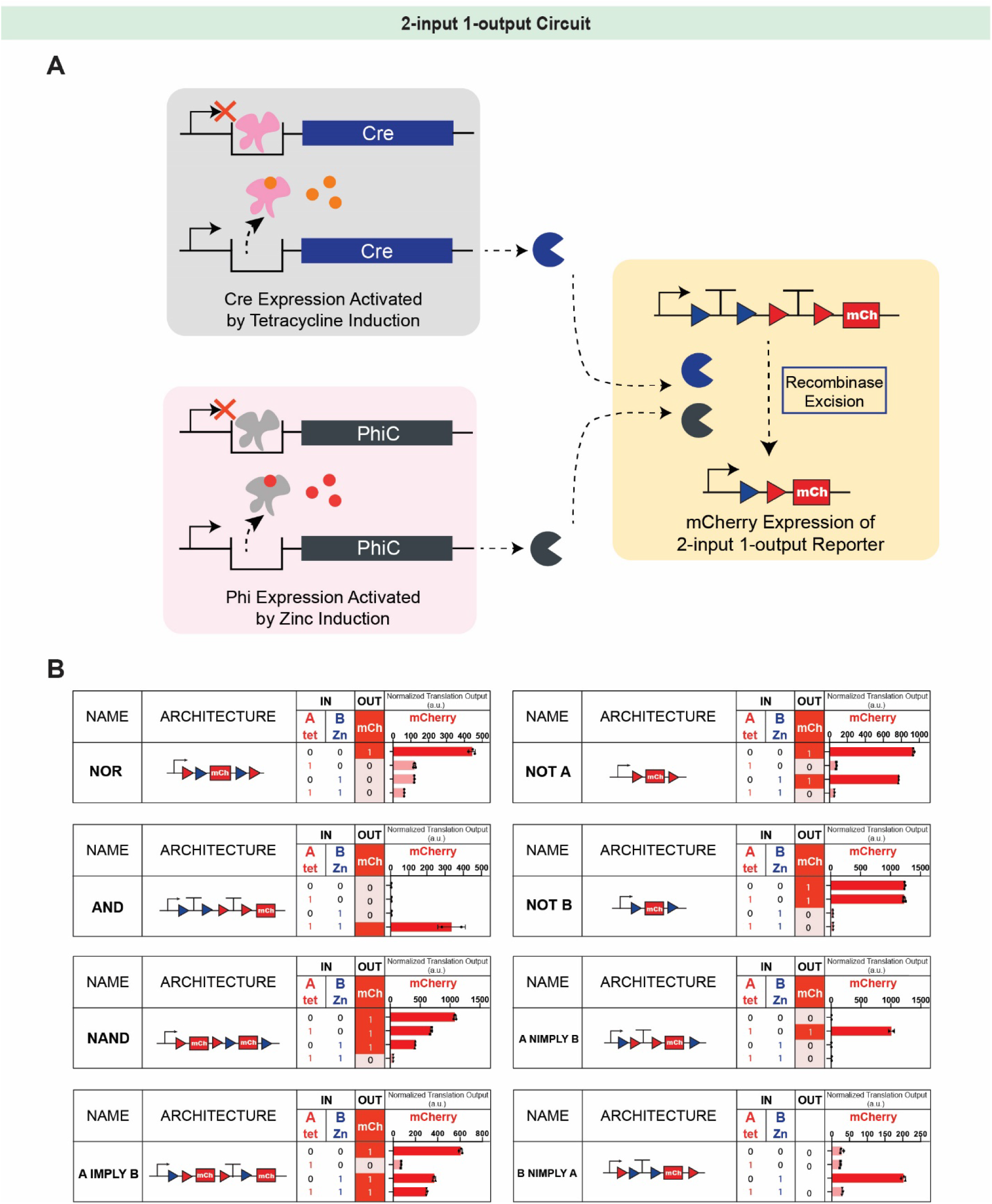
2-input-1-output circuits for two chemical sensing in the cell-free condition. (A) Schematic for the mechanism of the aTF-implemented 2-input-1-output circuit. Cre is controlled by the tetR system, while PhiC excision circuits are controlled by the smtB system. (B) Results of 2-input-1-output circuits for detecting tetracycline and zinc. Cre is controlled by tetR, while PhiC is controlled by smtB. To set up the experiment, recombinase plasmids, purified aTFs, water, and sensing molecules are added to the PURE reaction mix. After 30 minutes of incubation at 37°C, reporter plasmids were added to the reaction. Data shown are the means of technical triplicate or duplicate samples with error bars indicating +1 standard deviation. P-values were calculated as described in the methods.

The performance of the circuits was evaluated by a vector proximity (VP) metric measuring the discrepancy between the experimental output and the expected logic table. The expected output indicated by the logic table and the experimental measurements were represented by a truth table vector and a signal vector, respectively, in a four-dimensional space ^36^ (**Supplementary** Fig. 8A). The angular difference between the two vectors was measured: a 0° angle indicates that the experimental results represent the intended logic table perfectly, and a 90° angle indicates that the experimental output demonstrates completely wrong output. Among the 11 circuits tested, only 5 circuits (36%) showed a VP angle that was smaller than 15° using the original experimental procedure (**Supplementary** Fig. 8B). With the modified procedure, the circuits’ performance improved, and the VP angle of all circuits systematically decreased, resulting in 6 of 8 circuits (80%) with VP angle smaller than 15°. Additionally, 2-input 1-output circuits with biosensors yield 7 over 8 circuits (88%) with a VP angle smaller than 15°.

### A 2-Input-4-Output BLADE Circuit in Cell-free

Although 2-input-1-output circuits have successfully developed in cell-free systems^44^, a general strategy for creating N-input-M-output has yet to be developed. The Wong lab has previously developed a 2-input-4-ouptut ‘Boolean logic and arithmetic through DNA excision’ (BLADE) in mammalian cells to exploit the features of site-specific recombinases to enable N-input-M-output combinatorial computation^36^. With two inputs, the circuit requires four different outputs, Z = Z_AB_ = Z_00_, Z_10_, Z_01_, and Z_11_, to represent 2^N^ = 2^2^ = 4 (N indicates input number) different states of inputs A and B (**Fig. 5A**). To investigate the potential of building N-input-M-output in a cell-free environment, we implemented the 2-input BLADE circuits with terminators, fluorescent, and luminescent genes. One key element in the BLADE circuit is the heterospecific site, such as the Cre recombinase’s lox66/72 and lox2272. Although they are only a few base pairs different, lox66/72 and lox2272 can only retain Cre excision capability when paired with their corresponding recognition sites, which means lox2272 can only be paired with lox2272 to trigger Cre recombination activity while lox66 can only be paired with lox72. With constitutively expressed recombinases Cre and PhiC, reporter GFP, mCherry, and NanoLuc expression was induced under corresponding states (**Supplementary** Fig. 7). Furthermore, the BLADE circuits, when combined with aTFs-based biosensors, can be used for environmental surveillance for small molecules and heavy metal ions such as tetracycline and zinc (**Fig. 5B**). The VP angle metric for the BLADE circuits was evaluated to be less than 25%, for all fluorescent and luminescent outputs (**Supplementary Table 8B**).

**Figure 5.**
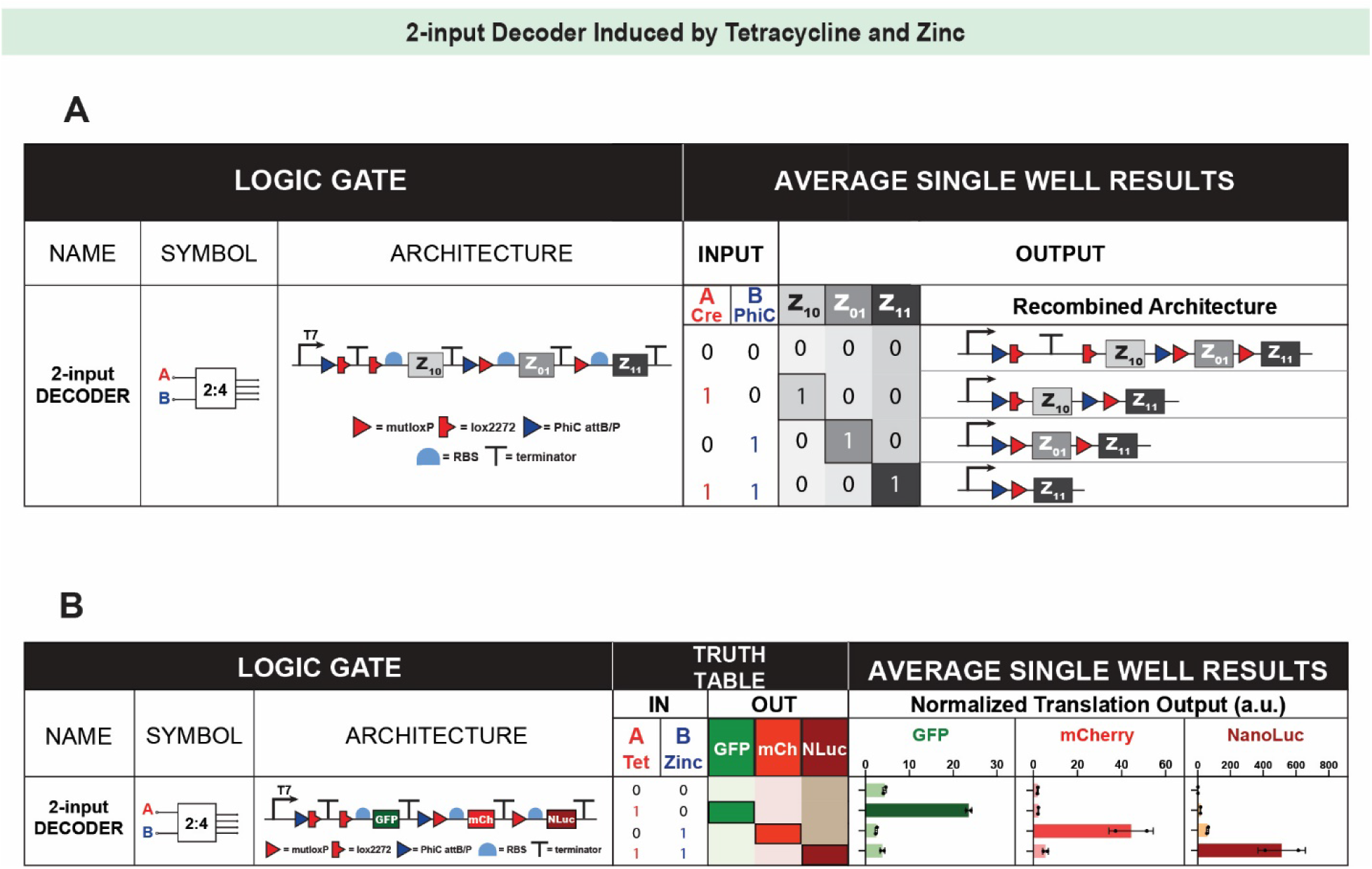
2-input-4-output Decoder. (A) Schematic for the mechanism of the 2-input decoder. The reporter plasmid expression cassette is separated into four addresses by recombinase recognition sites, resulting in 4 different outputs based on four states of two inputs. (B) Results of 2-input-4-output decoder for detecting tetracycline and zinc. Cre is controlled by the tetR system, while PhiC is controlled by the smtB system. To set up the experiment, recombinase plasmids, aTFs, water, and sensing molecules are mixed and added to the PURE reaction mix. After 30 minutes of incubation at 37°C, reporter plasmids were added to the reaction. Data shown are the means of technical duplicate samples with error bars indicating +1 standard deviation. P-values were calculated as described in the methods.

### Cell-Free Biological Memory Storage with CRIBOS

The capability to record crucial biological memory shapes research and extends to diverse applications in disease diagnostics and medical therapeutics. For example, immunological memory enables the development of innovative therapeutic interventions such as vaccines and immunotherapy^45,46^. Additionally, DNA barcoding plays a pivotal role in detecting invasive organisms and uncovering novel species^47–49^. These applications underscore the far-reaching impact of biological memory storage in advancing various fields of science and technology.

DNA has been used to transmit genetic information to the next generations because of its high stability, storage capacity, small volume, and inheritance. Moreover, employing DNA as the substrate for memory storage facilitates direct interfacing with sensors capable of detecting significant biological and environmental signals. This integration allows for the activation of biological programs in response to various environmental changes and bridges the gap between sensing and biological responses. Site-specific recombinase permanently modifies DNA sequence with high fidelity and efficiency, serving as a powerful writer to encode biological information into the DNA-based memory storage medium. Thus, with the success of building recombinase-based Boolean logic gates in a cell-free environment, we created a portable, robust, and stable biological memory storage device with the CRIBOS platform.

To evaluate the portability potential, we apply CRIBOS in a paper-based setting. One benefit of the PURE system is that it can be lyophilized on a sterile and abiotic paper-based platform. The lyophilized paper-based cell-free reaction is stable at room temperature for over a year and can be activated through rehydration^50,51^. In contrast, liquid-based reactions must be stored at −80°C to maintain enzyme activity.

In addition to device portability, memory stability is critical for a memory storage device. DNA memory can not only be effectively modified on paper through recombinase circuits but can also be stably preserved under room temperature for over four months upon activation (**Fig. 6B**). Paper-based CRIBOS with a plasmid reporter DNA and linear recombinase expressing gene was activated on day 0 through rehydration, and information was stored on the paper. At different time points post-activation, memory stored in the paper-based device was retrieved and analyzed. The reporter DNA with biological memory was retrieved through elution, and then collected DNA was used to transform bacteria to express mCherry reporter output (**Fig. 6A**). The recombinase excision rate was evaluated based on the percentage of mCherry+ bacteria colonies. High excision efficiency in the reporter+Cre condition indicates that the information was well-preserved at room temperature, and no DNA mutation is observed based on the sequencing results (**Fig. 6B and Supplementary** Fig. 9).

**Figure 6.**
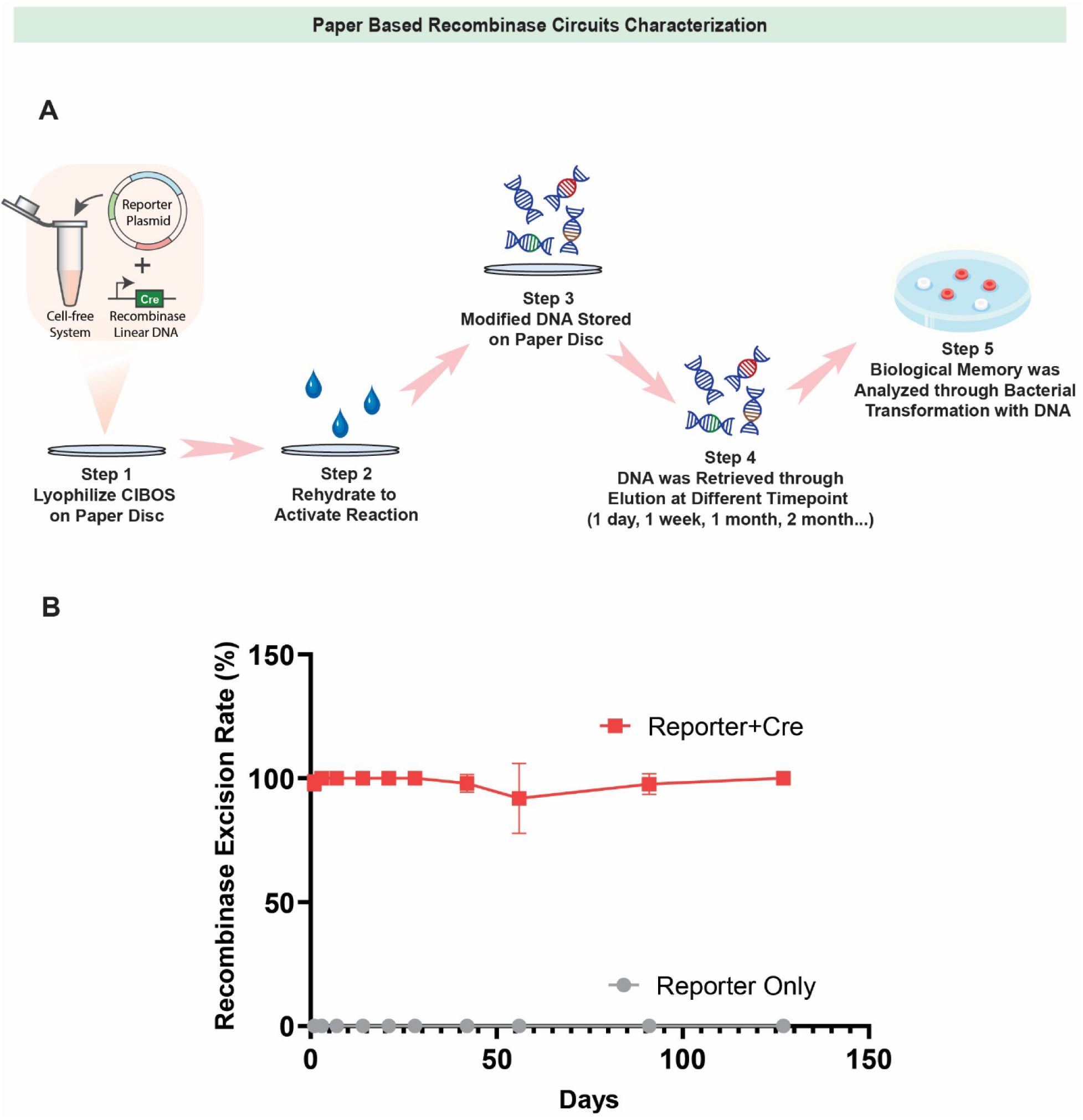
Paper-based recombinase circuits and memory recording platform. (A) Schematic of the experimental process for paper-based cell-free recombinase setup. Reporter plasmid and linear recombinase DNA are mixed with PURE reaction mix and then freeze-dried onto a paper disc for stable storage under room temperature. Then, the reactions are activated through rehydration. Modified DNA-based memory was stored on the paper disc for different lengths of duration, ranging from 1 day to 4 months. On the day of analysis, DNA-based memory is retrieved from a paper disc and transformed into bacteria for signal output expression. (B) Memory stability evaluation for paper-based memory storage devices. Paper-based CRIBOS was activated through rehydration on day 0. DNA-based memory was collected on different days up until four months. Data shown are the means of technical triplicate samples with error bars indicating +1 standard deviation. P-values were calculated as described in the methods.

## Discussion

We introduce CRIBOS, a recombinase-based multiplex genetic circuit platform in a cell-free environment. We used CRIBOS to develop the world’s first 2-input-4-output Boolean logic gate for chemical sensing in a cell-free system. Our results show that six different recombinases can function in cell-free conditions with high activity, with over a 20-fold difference between with and without recombinases. Our system also has high orthogonality, enabling the construction of complex genetic logic gates.

Through rigorous characterization, we discovered multiple design rules for recombinase-based Boolean logic gates in the cell-free environment. The challenges encountered during the optimization and troubleshooting process illustrate that applying genetic circuits developed in cells to cell-free is non-trivial. Our design rules, therefore, will facilitate future recombinase circuit developments. Although cell-free is a natural environment for circuit development and can provide benefits and convenience that cellular platforms and animal models cannot provide, it poses challenges and issues not present in other platforms. For instance, while living organisms can offer genetic circuits with continuous and durable energy sources and building blocks, a cell-free environment has limited means to regenerate high-energy compounds and starting materials needed for protein synthesis and circuit components. This illustrates that proper resource allocation is critical for cell-free operations. Also, cell-free systems are more diluted than the cell lysate, not just in the concentration of the transcription and translation machinery but also in the degree of macromolecular crowding^52–55^. Such differences in spatial density can influence the biochemical reaction rates and equilibria, changing the efficiency and competitive effect of protein-protein and protein-DNA binding. The biochemical characteristics of recombinase-recognition sites binding might also be affected in the cell-free reaction, as suggested by the complete shutoff of the loxP reporter signal by Cre recombinase due to the high Cre-loxP binding affinity post recombination. While lox66/72 sites were previously used for fixing the directionality of the recombination, few studies have reported its functions in improving circuit output signals in cells or animal models because the native Cre-loxP system can function properly and efficiently in those platforms, and there is no need to replace the loxP sites into lox66/72 sites. These discrepancies between cell and cell-free systems will be critical considerations for building complex genetic circuits and expanding the cell-free bio-computation toolkits.

The capability of cell-free recombinases to construct multi-input-multi-output circuits was also demonstrated with CRIBOS. Using two of the six orthogonal recombinases, the BLADE circuit that was previously developed in mammalian cells was also designed and optimized for best circuit output in a cell-free environment. By implementing aTFs-based small molecule sensors, the BLADE circuit can be induced by multiple chemical inducers in parallel, which is applicable for multiplex environmental sensing and water contaminant detections^22^. With the BLADE circuit as the proof-of-principle, more complex genetic circuits can also be generated with CRIBOS. Previously, we developed a framework for building multi-input-multi-output genetic circuits in mammalian cells. Using the 2-input BLADE design as a basic structure, genetic circuits with three or more inputs can be efficiently designed and implemented. Besides engineering circuits to detect the concurrence of multiple events, recombinase-based temporal circuits can also be designed to detect sequences of events, providing an extra layer of information. Combined with CRIBOS, such complex systems can be expanded to a cell-free environment to broaden the toolkits further.

After demonstrating that complex bio-computation can be achieved with recombinase genetic circuits in cell-free conditions, we applied the platform with the aTFs that can serve as water contaminant sensors. Although our system only showcases small molecule sensing with aTFs-based sensors as a demonstration, CRIBOS can also be combined with various other sensors to enable diverse applications. Robust and efficient cell-free recombinases act as reliable writers of the input-output system, paving a solid foundation for the CRIBOS system and allowing for flexible modifications of other circuit modules. In mammalian cells, recombinase genetic circuits are also compatible with chemical-induced dimerization (CID)-based sensors^56^. Adapting cell-free recombinase genetic circuits with CID-based sensors can facilitate CRIBOS with the capability to detect changes in temperature, fluorescence, luminescence, and the presence of a broader range of chemicals.

Lastly, we demonstrated that CRIBOS can be applied in a paper-based setting to improve the portability and stability of the genetic circuit platform. Paper-based CRIBOS is the world’s first genetic circuit with memory lasting over four months with minimal resource, energy cost, and maintenance required. Methods to encode, store, and retrieve memory from the paper-based CRIBOS were explored and developed at room temperature and 37°C to demonstrate the broad application conditions of the memory storage device. While the current method for memory retrieval is cost- and resource-effective, more sensitive and rapid methods for detection out in the field can also be explored in future studies by combining with next-generation sequencing and genetic engineering tools. CRIBOS memory devices can also be tested and characterized under more diverse reaction conditions, such as under extremely high or low temperatures. In addition, although we only applied CRIBOS in a paper-based setting, cell-free reactions can function in various settings, such as microfluidics and flow strips. Thus, in the future, we can combine CRIBOS with microfluidics devices to further generate a portable memory storage device that allows for continuous and multiplex environmental sensing under a low-resource setting.

## Methods

### Strains and growth medium

NEB 5-alpha Competent E. coli (New England Biolabs, C2987U) was used for routine molecular cloning. E. coli strain Rosetta 2 (DE3) pLysS (Novagen, 71401) was used for recombinant protein expression. BL21(DE3) Competent *E. coli* (New England Biolabs, C2527H) was used for recombinase circuit testing. LB broth (Thermo Fisher Scientific, BP1426-2) supplemented with appropriate antibiotic(s) (100 μg/ml carbenicillin (Thermo Fisher Scientific, 10177012), 50 μg/ml kanamycin (Bio Basic, KB0286-25) and/or 25 μg/ml chloramphenicol (Sigma-Aldrich, C0378-25G)) was used as a bacterial growth medium. Bacterial transformations were performed based on the manufacturer’s protocols.

### Molecular cloning

DNA oligonucleotides for cloning and sequencing were synthesized by ThermoFisher Scientific. Genes encoding fluorescent, luminescent proteins and 2X terminators were synthesized as gBlocks (Integrated DNA Technologies). All constructs were created using standard restriction enzyme digestion, Gibson isothermal assembly, and Unique Nucleotide Sequence (UNS) Guided assembly procedures into a pET15b plasmid. UNS Guided assembly utilizes a mechanism similar to Gibson’s isothermal assembly but with standard homology sequences that were computationally optimized. Constructed plasmids were transformed and maintained in NEB 5-alpha (New England Biolabs (NEB) C2987H) *Escherichia coli* competent cells at 37°C overnight before miniprep (Epoch Life Sciences, 2160250).

### Protein Production and Purification

All aTFs expression plasmids were purchased from Addgene. For aTFs production, plasmids were transformed into Rosetta 2 (DE3) pLysS E.coli strain. A single colony of transformed bacteria was cultured in liquid LB media overnight. The next day, overnight cultures were diluted 1:100 into 200mL-500mL culture media. OD of culture media was measured by a plate reader every 30 minutes. When OD reached 0.5-0.8, culture was induced with 100uM Isopropyl β-D-thiogalactoside (IPTG) (Sigma-Aldrich, I6758-5G) overnight at 30°C. After overnight IPTG induction, bacterial culture was pelleted through a centrifuge at 3500Xg for 20 minutes. Every 1g of bacterial pellet is resuspended in 10ml SoluLyse lysis buffer (Amsbio, L200500) supplemented with 1 tablet of protease inhibitor cocktail (Sigma-Aldrich, 11836170001), 40mM imidazole (Fisher Scientific, O3196-500), 200uL Lysozyme (Sigma-Aldrich, L4919-500MG) and 0.5uL Benzonase (EMD Millipore, 71205-3). Lysozyme was pre-resuspended to 10mg/mL in water and stored at −80°C upon arrival. The resuspended cell pellet was placed on a mixer for 20 minutes and centrifuged at 4000g for 10 minutes at 4°C to remove bacterial debris post-incubation. The bacterial supernatant was then sterilely filtered with a 0.22µm filter (EMD Millipore, SCGP00525) to remove bacterial precipitate and avoid clogging in FPLC columns.

Clarified supernatants were purified using His-tag affinity chromatography with a HisTrap column (Cytiva, 17-0406-01), followed by Heparin chromatography with a heparin column (Cytiva, 17-5247-01) using an AKTAxpress fast protein liquid chromatography system. The fractions collected from the FPLC were buffer-exchanged and concentrated utilizing a protein concentrator (Thermo Scientific, 88515). Protein concentrations were evaluated using Nanodrop. The purity and size of proteins were validated with SDS-PAGE and western blot. Purified proteins were stored in PBS at 4°C.

### In vitro protein synthesis

NEB PURExpress In Vitro Protein Synthesis Kit (New England Biolabs, E6800S) supplemented with RNase Inhibitor (Sigma-Aldrich, 03335402001) was used to generate data for cell-free experiments based on the manufacturer’s protocol. The concentrations of plasmids and linear DNAs are listed in the Supplement. The plasmids, DNAs, aTFs, and small molecules are first premixed before combining with the cell-free reaction mix. Then, 5μL of cell-free circuits were incubated in each well of a 384-well clear bottom black plate (Corning, 3542) at 37°C for 5hr. An endpoint measurement, or dynamic measurement, was performed on a plate reader (SpectraMax M5, Molecular Devices) with an excitation wavelength of 580nm and emission wavelength of 611nm for mCherry, an excitation wavelength of 485nm and emission wavelength of 525nm for GFP, and an excitation wavelength of 381nm and emission wavelength of 445nm for BFP. Nano-Glo Luciferase Assay (Promega, N1110) was used for NLuc luminescence detection. Nano-Glo Luciferase Assay Substrate was diluted 1:50 by Nano-Flo Luciferase Assay Buffer. The luminescence level was measured immediately after a volume of the diluted substrate equal to 6X the volume of the cell-free reaction was added to each well.

A linear DNA template with a T7 promoter, followed by a recombinase-expressing gene and a T7 terminator, was PCR (CoWin Biosciences CW2965F) amplified from the corresponding recombinase plasmids using primers listed in the Supplement. Amplified DNA was purified with a PCR cleanup kit (Epoch Life Science 2360250), and band size and sequence were verified through gel electrophoresis and sequencing.

### Freeze-drying

Filter paper (Cytiva, 1442-070) for depositing and freeze-drying cell-free reactions was first treated with 5% bovine serum albumin (Sigma-Aldrich, A3059-100G) overnight at 37°C. The paper was then cut into 2-mm-diameter paper discs with a biopsy punch and placed at the bottom of a 96-well PCR plate. Before lyophilization, 2μL of cell-free reaction mix was applied directly onto the paper disc, and the PCR plate was sealed by porous AeraSeal Sealing Films (Excel Scientific, B-100). Post-freezing by liquid nitrogen, the PCR plate was immediately sent for lyophilization overnight. After lyophilization, the paper discs were inserted into the bottom of the 384-well plate and activated by adding 2μL of autoclaved DI water.

### Plate reader quantification and MEF standardization

A Texas Red Dye (Sigma-Aldrich, F6377-100G) was used to convert arbitrary mCherry fluorescence intensity into nanomolar equivalent Texas Red Dye (labeled as normalized translational expression). A serial dilution was prepared with PBS from a 500uM stock solution. The samples were prepared with triplicates, and fluorescence values were measured at an excitation wavelength of 580nm and emission wavelength of 611nm on a plate reader (SpectraMax M5, Molecular Devices). Fluorescence for a concentration in which a single replicate saturated at the plate reader was excluded from the analysis. The rest of the measurements were averaged and formed a linear regression line, which is used to convert the measured arbitrary fluorescence value to the concentration of the Texas Red Dye (**Supplementary** Fig. 7). Standard curves were created with the same process for each plate reader for data collection to normalize the results. The same process was used to convert GFP fluorescence intensity into nanomolar equivalent Fluorescein (Sigma-Aldrich, F6377) and NanoLuc luminescence intensity into nanomolar equivalent NanoLuc purified protein (Promega, E499A). All fluorescence and luminescence outputs are normalized to the nanomolar equivalent of corresponding dye or proteins and then present on the figure as “Normalized Translational Output (NTR).”

### Diagnostic analysis

For each inducer and recombinase combination, the limit of blank (LoB) was calculated as the mean of the blank added to three times the standard deviation of the blank. The limit of detection (LoD) was calculated as the LoB plus three times the standard deviation of low concentration sample. The dynamic range was computed as the mean expression of highest output for tetracycline and zinc positive samples over the mean expression of negative samples. False positive rate and true positive rate were used to construct the receiver operating characteristic (ROC) curve and compute the area under the curve (AUC).

### Vector proximity computational analysis

Each 2-input-1-output or 2-input-4-output Boolean function corresponds to a logic table with four rows and one output column or four rows and four output columns, correspondingly, the output column in each Boolean function was mapped to a 4-dimensional vector ***a***, called the “truth table vector”. The fluorescent reporter output signal of each genetic circuit measured from experimental implementation was also mapped to a 4-dimensional vector ***b***, called the “signal vector.” We evaluated the accuracy of a genetic circuit by computing the angle ***θ*** between the vector ***a*** and ***b*** using the formula:

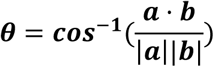

When computing ***θ***, we capped the signal values in ***b*** to a maximum of 400nM Texas Red, 60nM Fluorescein, and 400nM NanoLuc purified protein. The angular difference ranges from 0° (best) to 90° (worst).

### Statistical analysis

All cell-free in vitro experiments involved setting up PURE in vitro protein synthesis expression of DNA into *n* = 2 or 3 reactions. Fluorescence and luminescence intensities for each reaction were averaged, and the standard deviation was taken. To evaluate statistical significance, data between groups were evaluated with unpaired two-tailed t-tests. P values are reported (not significant = p > 0.05, * = 0.01 < p < 0.05, ** = 0.001 < p < 0.01, ∗∗∗ = 0.01 < p < 0.001, **** = 0.001 < p < 0.0001).

## Abbreviation

CRIBOS: Cell-Free Recombinase Integrated Boolean Output System

PURE: Protein Synthesis Using Recombinant Elements

BLADE: Boolean Logic and Arithmetic through DNA Excision

aTFs: Allosteric Transcription Factors

## Acknowledgment

W.W.W and D.D. acknowledge funding from the National Science Foundation (NSF) SemiSynBio-II (award No. 2027045). We also thank Wong lab members for suggestions on the manuscript; BU BME core facility for lyophilization device maintenance; K. Wu (Boston University), Z. Yan (Boston University), and A. A. Green for helpful discussion on the PURE system and paper-based cell-free reaction; P. Garden (Boston University) for helpful discussion on protein purification; N. Tague (Boston University) and M. Sheets (Boston University) for plasmids and materials for bacterial recombinase circuits; J. B. Lucks (Northwestern University) and J. Li for helpful discussion and testing samples of purified aTFs proteins; A. Wang, J. Ali, A. Fecteau and A. Tillisch for assistance in molecular cloning. Diana Arguijo (Boston University) for helpful discussions about the transition of our systems to continuous flow microfluidics.

## AUTHOR CONTRIBUTIONS

J.C. made molecular and cellular reagents, performed experiments, and generated all figures. J.C. and A.B. analyzed data. J.C., A.B., D.D., and W.W.W. wrote the paper.

## SUPPLEMENTAL INFORMATION

Document S1. Figures S1-S10 and Table S1-S24

