## Supplemental Figure for "Cell-Free Recombinase Integrated Boolean Output System (CRIBOS)"

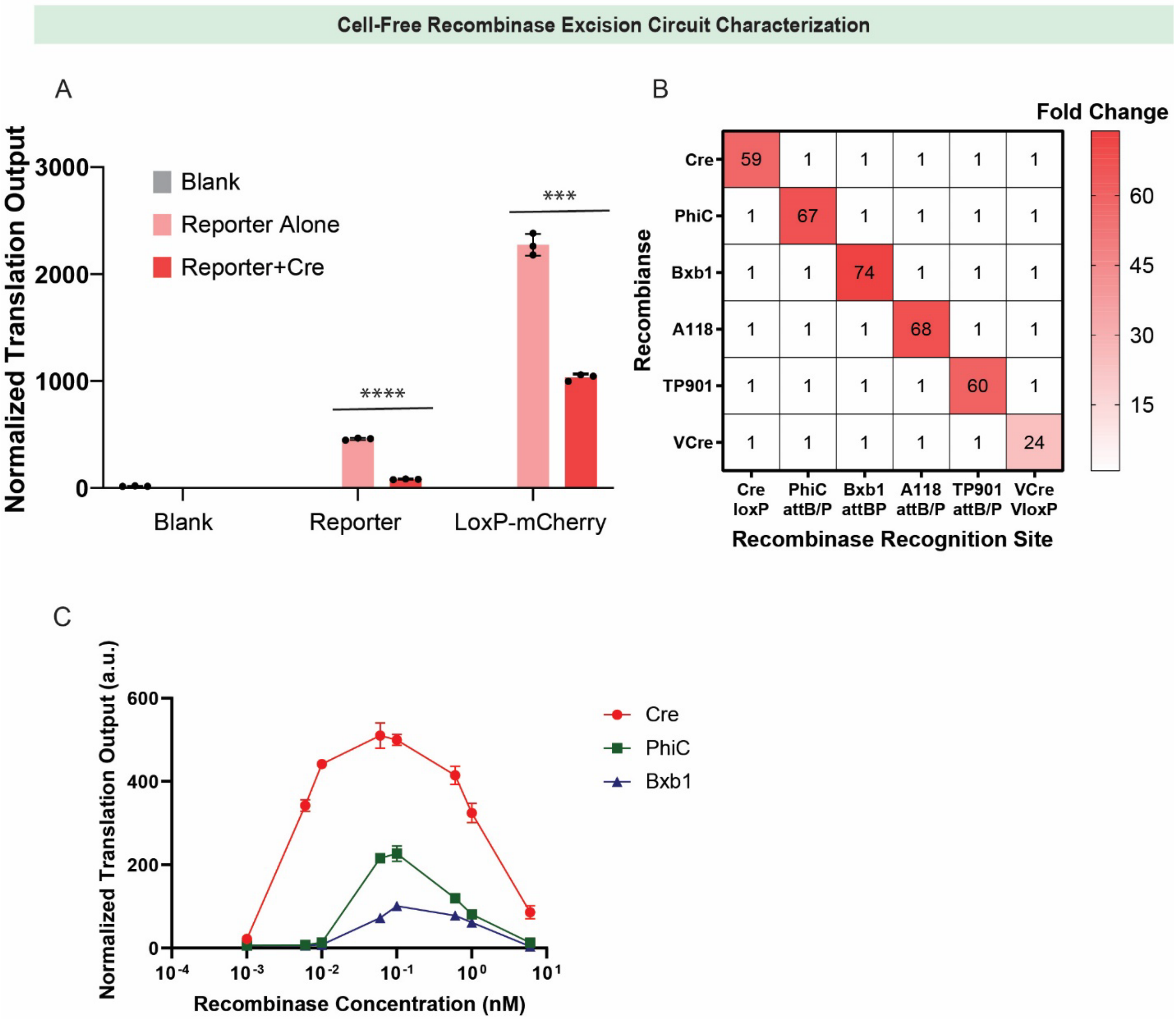


**Supplementary Figure 1:**

(A) The original Cre excision reporter is compared with blank negative control (water and cell-free reaction mix only) and loxP-mCherry plasmid.

(B) Recombinases are tested in the cell-free environment for their efficiency and orthogonality.

(C) The dose response for recombinase plasmid concentration is tested to optimize circuit function.

Data shown are the means of technical triplicate samples with error bars indicating +1 standard deviation. P-values were calculated as described in the methods.

Design Rules

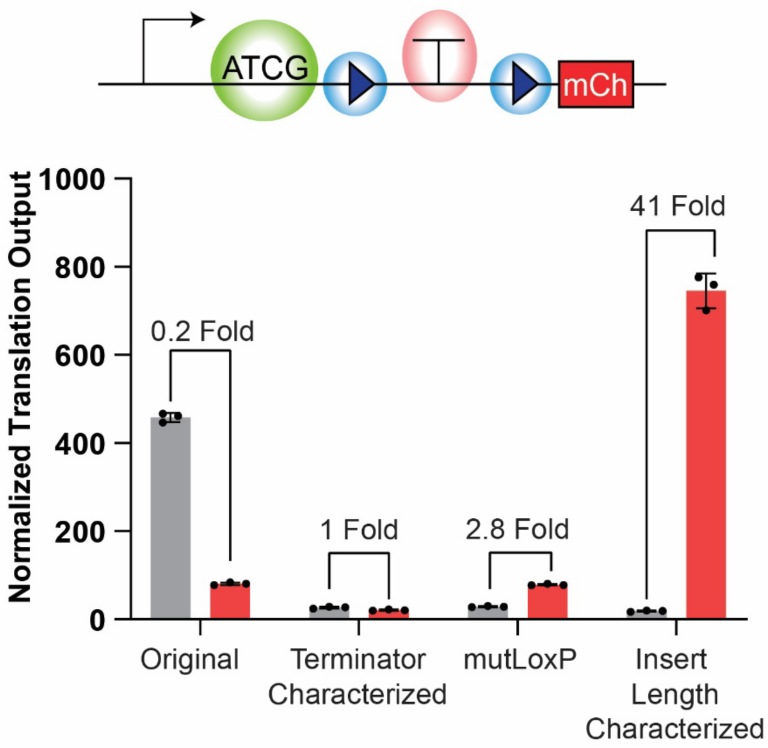


**Supplementary Figure 2:**

Characterization of excision reporter. Three sites on the reporter were optimized to improve circuit output: terminator sequence (red circle), recognition sites (blue circle), and insert length between promoter and terminator (green circle).

Data shown are the means of technical triplicate samples with error bars indicating +1 standard deviation. P-values were calculated as described in the methods.

**TF-controlled Recombinase Genetic Circuits**

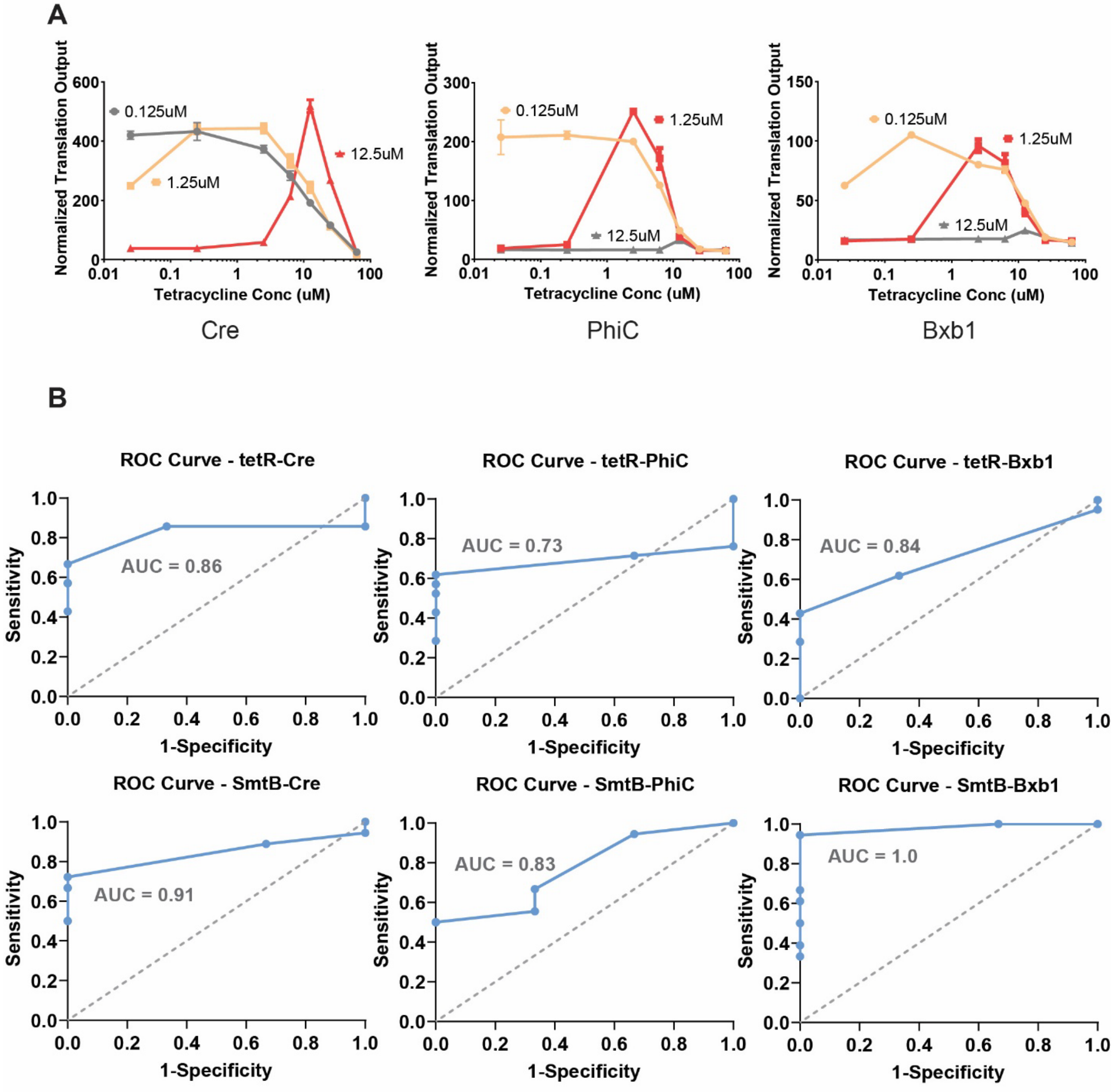


**C**

| tetR-tetracycline Biosensor | | | smtB-Zinc Biosensor | | |
|---|---|---|---|---|---|
| Recombinase | Detection Window (uM) | Dynamic Range of Sensing (uM) | Recombinase | Limit of Detection (uM) | Dynamic Range of Sensing (uM) |
| Cre | 1.08 – 59.0 | 14.0 | Cre | 1.17 | 20.7 |
| PhiC | 0.254 – 19.6 | 14.6 | PhiC | 2.69 | 8.14 |
| Bxb1 | 0.366 – 22.5 | 5.76 | Bxb1 | 4.45 | 12.8 |

**Supplementary Figure 3:**

(A) Dose-response curves of tetracycline measured by Cre, PhiC, and Bxb1 excision circuits using different concentrations of tetR proteins.

(B) ROC curves (receiver operating characteristic curve) and AUC (Area Under the ROC Curve) for aTFs-controlled recombinase excision circuits.

(C) Limit of detection, detection window, and dynamic range of sensing for aTFs-controlled recombinase excision circuits.

Data shown are the means of technical triplicate samples with error bars indicating +1 standard deviation. P-values were calculated as described in the methods.

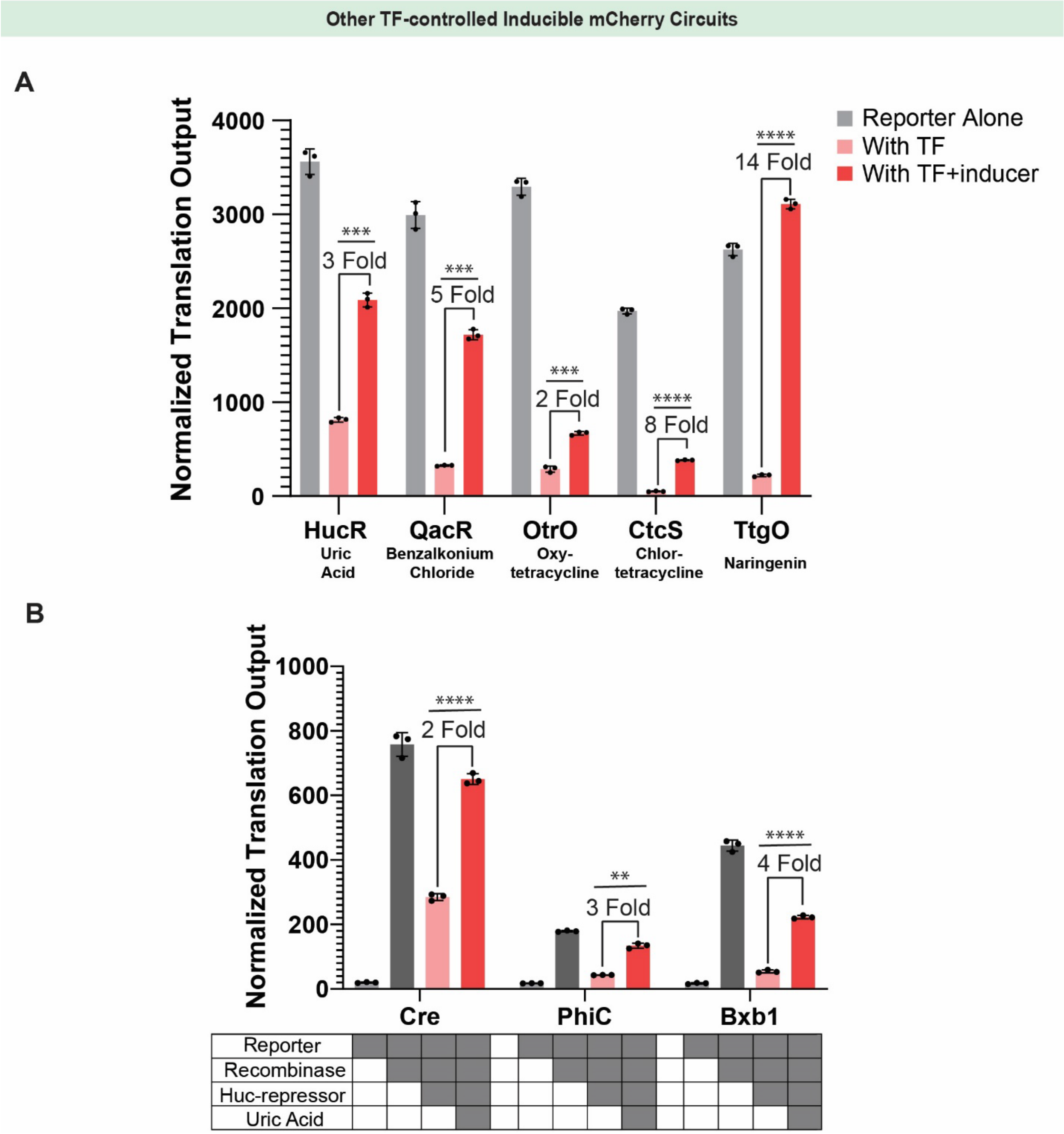


**Supplementary Figure 4:**

(A) Potential aTFs systems to control constitutive mCherry expression in the cell-free system.

(B) Cre, PhiC, and Bxb1 excision circuits controlled by HucR-uric acid aTFs.

Data shown are the means of technical triplicate samples with error bars indicating +1 standard deviation. P-values were calculated as described in the methods.

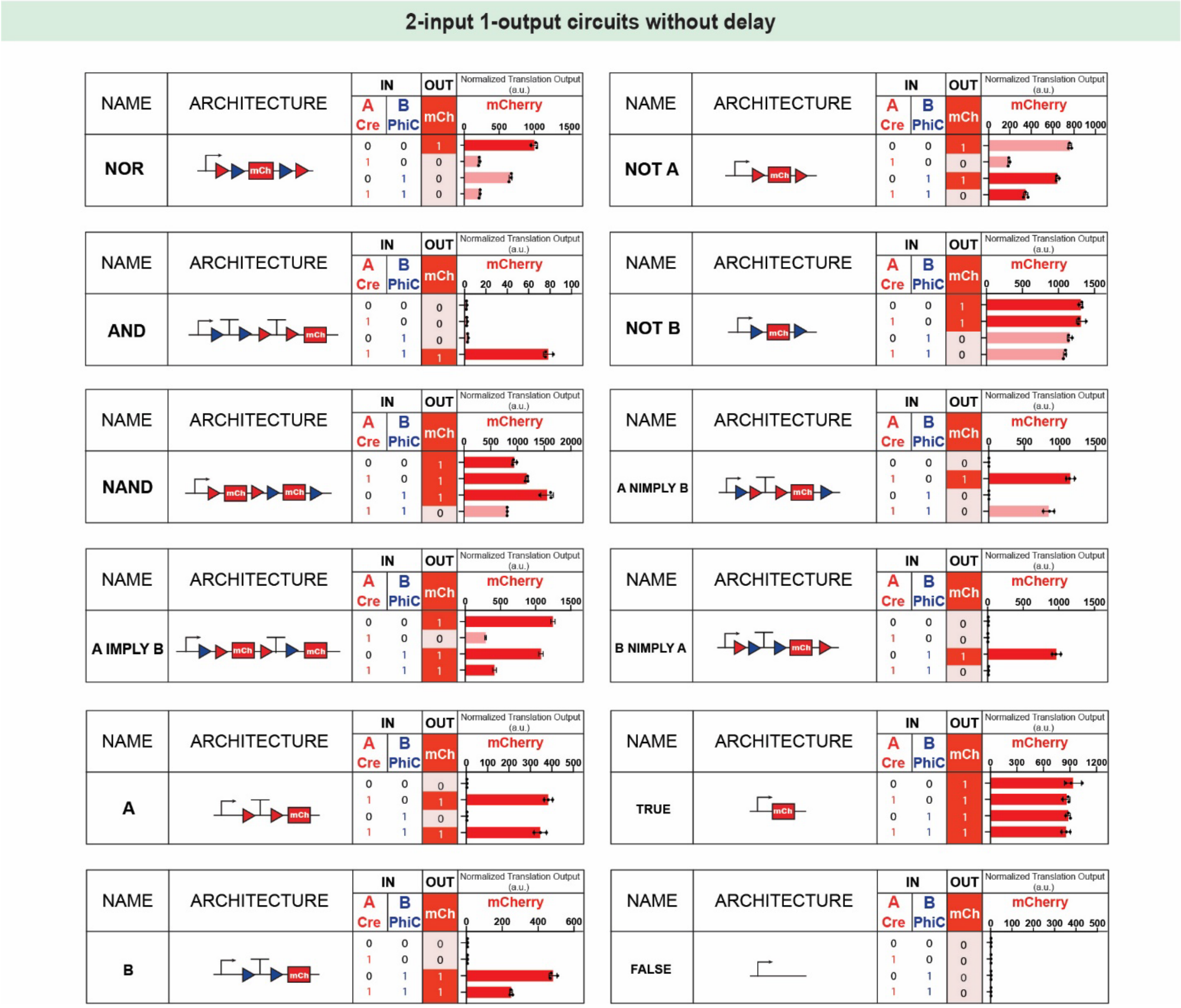


**Supplementary Figure 5:** Results of 2-input-1-output genetic circuits.

Data shown are the means of technical triplicate samples with error bars indicating +1 standard deviation. P-values were calculated as described in the methods.

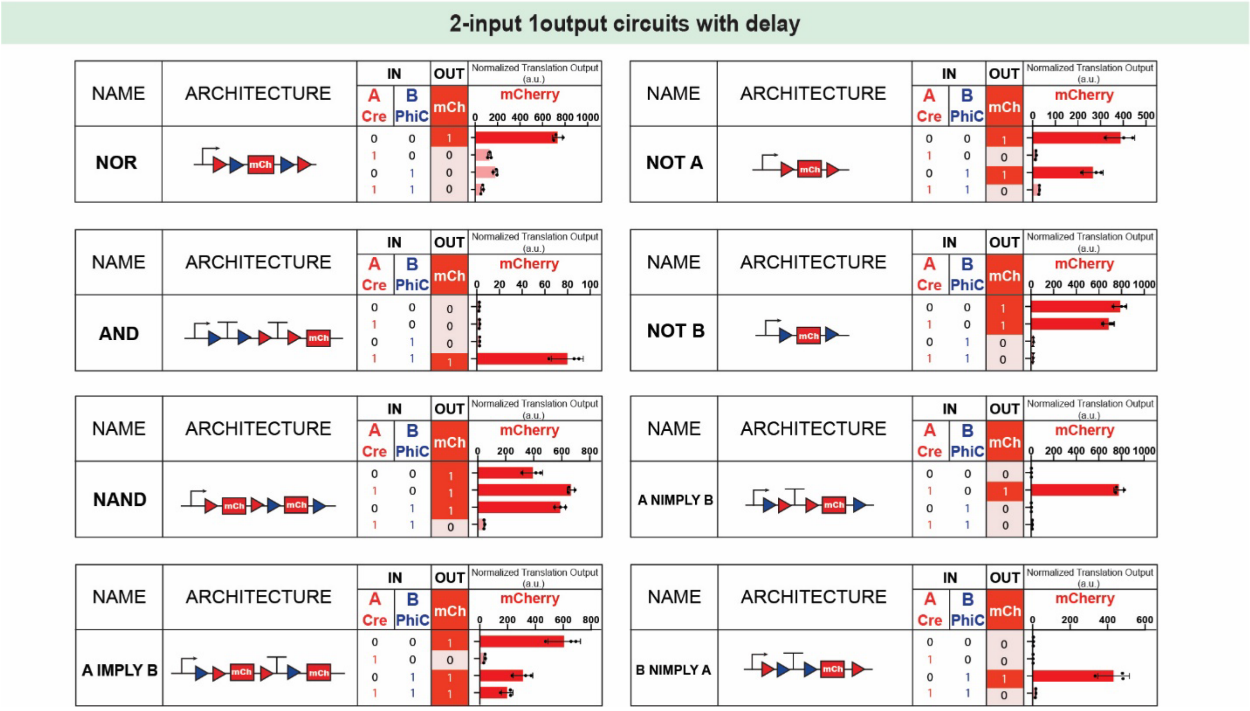


**Supplementary Figure 6:** Results of 2-input-1-output genetic circuits with improved experimental protocol. Recombinase plasmids were added to the PURE reaction mix to initiate the reaction. After 30 minutes of incubation at 37°C, reporter plasmids were added to the reaction.

Data shown are the means of technical triplicate samples with error bars indicating +1 standard deviation. P-values were calculated as described in the methods.

2-input 4-output Decoder

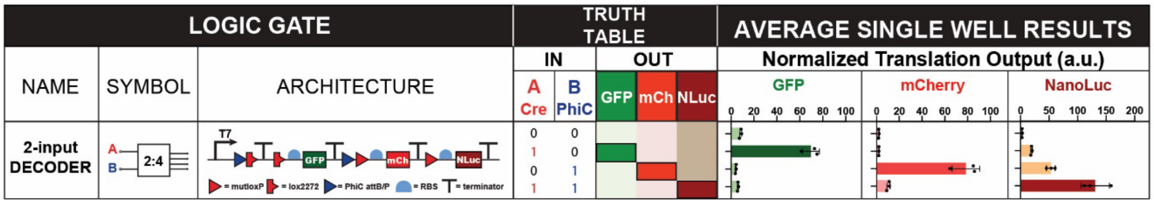


**Supplementary Figure 7:** Results of 2-input-4-output decoder. GFP, mCherry, and NanoLuc arbitrary measurements are all normalized to equivalent nanomolar of Fluorescein, Texas Red, and purified NanoLuc proteins based on the standard curves. The dye or protein concentration corresponds to the arbitrary measurements present in the figure as “Normalized Translation Output.”

Data shown are the means of technical triplicate samples with error bars indicating +1 standard deviation. P-values were calculated as described in the methods.

**Vector Proximity Metric Analysis**

**A**

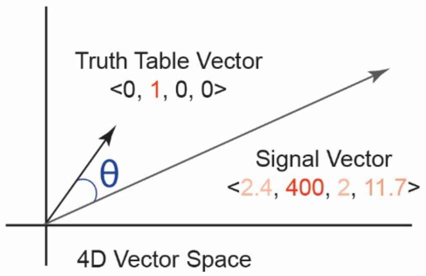


**B**

| 2 Input 1 Output Circuits Without Delay | | | |
|---|---|---|---|
| Circuit Type | VP Angle | Circuit Type | VP Angle |
| TRUE | 0 | NAND | 30.0 |
| B NIMPLY A | 2.5 | NOT A | 35.7 |
| A | 2.9 | A NIMPLY B | 45.0 |
| AND | 3.2 | NOT B | 45.0 |
| B | 13.1 | NOR | 51.7 |
| A IMPLY B | 22.7 | | |

| 2 Input 1 Output Circuits With Delay | | | |
|---|---|---|---|
| Circuit Type | VP Angle | Circuit Type | VP Angle |
| A NIMPLY B | 1.7 | NAND | 4.4 |
| B NIMPLY A | 2.6 | NOT A | 10.1 |
| AND | 2.9 | A IMPLY B | 15.8 |
| NOT B | 2.9 | NOR | 30.0 |

| 2 Input 1 Output Circuits With aTFs-based Biosensors | | | |
|---|---|---|---|
| Circuit Type | VP Angle | Circuit Type | VP Angle |
| AND | 1.3 | A IMPLY B | 9.2 |
| A NIMPLY B | 1.7 | NOT A | 9.3 |
| NAND | 4.1 | B NIMPLY A | 13.3 |
| NOT B | 5.7 | NOR | 24.3 |

| 2 Input 4 Output Decoder | | | | | |
|---|---|---|---|---|---|
| Output | VP Angle | Output | VP Angle | Output | VP Angle |
| GFP | 15.1 | mCherry | 8.1 | NLuc | 9.0 |
| 2 Input 4 Output Decoder with aTFs-based Biosensors | | | | | |
| Output | VP Angle | Output | VP Angle | Output | VP Angle |
| GFP | 10.5 | mCherry | 7.8 | NLuc | 23.4 |

**Supplementary Figure 8: Vector proximity metric analysis**

(A) The truth table of each genetic circuit is mapped into a four-dimensional vector called a “Truth Table Vector.” The experimental measurement of each genetic circuit is also mapped into a four-dimensional vector called a “Signal Vector.” To evaluate the correctness of the genetic circuit, the angle θ between the Truth Table Vector and Signal Vector is calculated for vector proximity (VP) metric analysis. We defined the circuits with VP angle <15° as functioning with high robustness.

(B) VP angles of 2-input-1-output and 2-input-4-output genetic circuits.

**Paper-Lyophilized Recombinase Circuit Characterization**

>> Reporter Alone sequence:
TAATACGACTCACTATAGGAGCGGCCGCACTAGTTACCGTTCGTATAGCATACATT
ATACGAAGTTATCCAGGCATCAAATAAGGATCCAAACTCGAGTAAGGATCTCCTAG
CATAACCCCGCGGGGCCTCTTCGGGGGTCTCGCGGGGTTTTTTGCTGAAAGAA
GCTTCAAATAAAACGAAAGGCTCAGTCGAAAGACTGGGCCTTTCGTTTTATCTGT
TGTTTGTCGCTGCGCGGCCGCCCTAGCATAACCCCTTGGGGCCTCTAAACGGGT
CTTGAGGGGTTTTTTGGTCGACCTAGCATAACCCCGCGGGGCCTCTTCGGGGGT
CTCGCGGGGTTTTTTGCTGAAAGAAGCTTCAAATAAAACGAAAGGCTCAGTCGAA
AGACTGGGCCTTTCGTTTTATCTGTTGTTTGTCGCTGCGCGGCCGCCCTAGCATA
ACCCCTTGGGGCCTCTAAACGGGTCTTGAGGGGTTTTTTGCCATAACTTCGTATA
GCATACATTATACGAACGGTACCTCTAGAAATAATTTTGTTTAACTTTAAGAAGGAGA

>> Reporter+Cre White Colony sequence:
TAATACGACTCACTATAGGAGCGGCCGCACTAGTTACCGTTCGTATAGCATACATT
ATACGAAGTTATCCAGGCATCAAATAAGGATCCAAACTCGAGTAAGGATCTCCTAG
CATAACCCCGCGGGGCCTCTTCGGGGGTCTCGCGGGGTTTTTTGCTGAAAGAA
GCTTCAAATAAAACGAAAGGCTCAGTCGAAAGACTGGGCCTTTCGTTTTATCTGT
TGTTTGTCGCTGCGCGGCCGCCCTAGCATAACCCCTTGGGGCCTCTAAACGGGT
CTTGAGGGGTTTTTTGGTCGACCTAGCATAACCCCGCGGGGCCTCTTCGGGGGT
CTCGCGGGGTTTTTTGCTGAAAGAAGCTTCAAATAAAACGAAAGGCTCAGTCGAA
AGACTGGGCCTTTCGTTTTATCTGTTGTTTGTCGCTGCGCGGCCGCCCTAGCATA
ACCCCTTGGGGCCTCTAAACGGGTCTTGAGGGGTTTTTTGCCATAACTTCGTATA
GCATACATTATACGAACGGTACCTCTAGAAATAATTTTGTTTAACTTTAAGAAGGAGA

>> Reporter+Cre Red Colony sequence:
TAATACGACTCACTATAGGAGCGGCCGCACTAGTTACCGTTCGTATAGCATACATTA
TACGAACGGTACCTCTAGAAATAATTTTGTTTAACTTTAAGAAGGAGA

**Supplementary Figure 9: Paper-lyophilized recombinase circuit characterization**

DNA-based memory validation through sequencing. DNA eluted from paper discs was used to transform bacteria, and colonies were picked. The three representative examples shown here are from a reporter (white colony) only condition and a reporter+Cre (white and red) condition. The T7 promoter is indicated by green, Cre recognition sites are blue, terminators are red, and the ribosome binding site is yellow.

**Standard Curve**

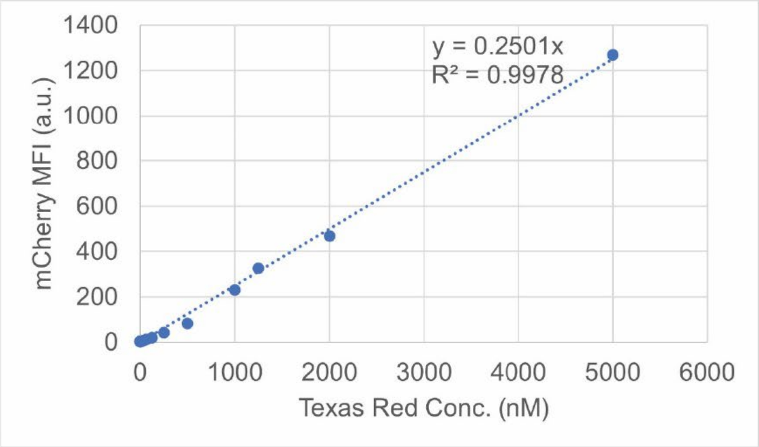


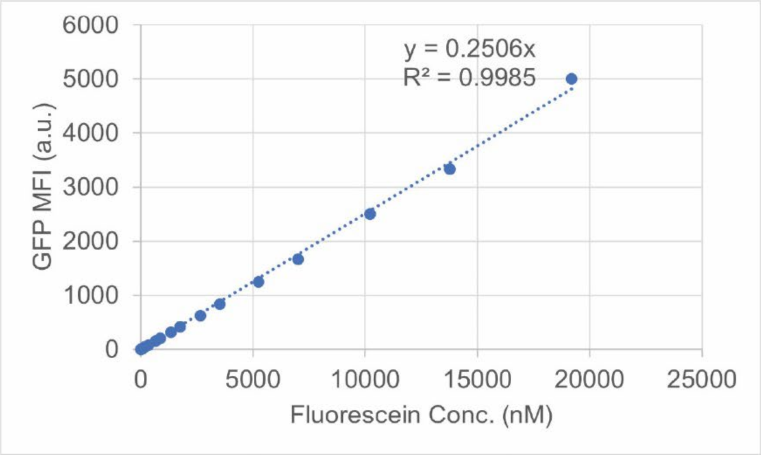


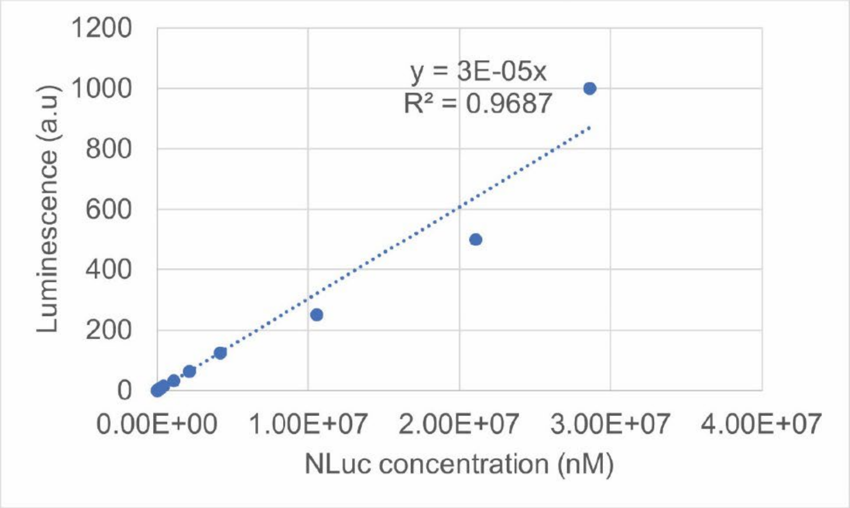

**Supplementary Figure 10: Standard curves for fluorescence and luminescence expression.**

Arbitrary Units of fluorescence and luminescence were standardized to nM concentration of Texas Red, Fluorescein (FITC), and NanoLuc. In the three representative examples shown here, the Texas Red and FITC dilution series was prepared with PBS, and NanoLuc was prepared in Promega Nano-Glo Luciferase Assay Buffer (Promega, E499A). The dilution series was measured on a plate reader using the same settings for measuring mCherry, GFP, and NanoLuc from PURE reactions. Data shown are for n=3 replicates.

Supplementary Table 1: Concentration of DNA for cell-free reaction in Figure 2B

| Reporter | Plasmid Number | Plasmid Conc. (nM) |
|---|---|---|
| Terminator 1 | JC42 | 6 |
| Terminator 2 | JC48 | 6 |
| Terminator 3 | JC464 | 6 |
| Terminator 4 | JC465 | 6 |

Supplementary Table 2: Concentration of DNA for cell-free reaction in Figure 2C

| Reporter | Plasmid Number | Plasmid Conc. (nM) | Recombinase | Plasmid Number | Plasmid Conc. (nM) |
|---|---|---|---|---|---|
| loxP-mCherry | JC466 | 6 | Cre | JC40 | 0.6 |
| mutLoxP-mCherry | JC467 | 6 | | | |

Supplementary Table 3: Concentration of DNA for cell-free reaction in Figure 2D

| Reporter | Plasmid Number | Plasmid Conc. (nM) | Recombinase | Plasmid Number | Plasmid Conc. (nM) |
|---|---|---|---|---|---|
| 0bp | JC757 | 6 | Cre | JC40 | 0.6 |
| 5bp | JC758 | 6 | | | |
| 10bp | JC759 | 6 | | | |
| 15bp | JC213 | 6 | | | |
| 20bp | JC760 | 6 | | | |
| 30bp | JC761 | 6 | | | |

Supplementary Table 4: Concentration of DNA for cell-free reaction in Figure 2E

| Reporter | Plasmid Number | Plasmid Conc. (nM) | Recombinase | Plasmid Number | Plasmid Conc. (nM) |
|---|---|---|---|---|---|
| Original | JC42 | 6 | Cre | JC40 | 0.6 |
| Modified | JC213 | 6 | | | |

Supplementary Table 5: Concentration of DNA for cell-free reaction in Figure 3B

| Reporter | Plasmid Number | Plasmid Conc. (nM) |
|---|---|---|
| tetO-mCherry | JC265 | 6 |
| smtO-mCherry | JC266 | 6 |

Supplementary Table 6: Concentration of proteins and chemicals for cell-free reaction in Figure 3B

| aTFs Protein | aTFs Protein Number | aTFs Protein Conc. (uM) | Chemicals | Chemicals Conc. (nM) |
|---|---|---|---|---|
| tetR | JC20 | 1.25 | tetracycline | 12.5 |
| smtB | JC22 | 5 | zinc | 12.5 |

Supplementary Table 7: Concentration of DNA for cell-free reaction in Figure 3C and 3D

| Reporter | Plasmid Number | Plasmid Conc. (nM) | Recombinase | Plasmid Number | Plasmid Conc. (nM) |
|---|---|---|---|---|---|
| Cre | JC213 | 6 | tetO-Cre | JC287 | 0.6 |
| PhiC | JC196 | 6 | tetO-PhiC | JC288 | 0.6 |
| Bxb1 | JC271 | 6 | tetO-Bxb1 | JC289 | 0.6 |
| | | | smtO-Cre | JC603 | 0.6 |
| | | | smtO-PhiC | JC604 | 0.6 |
| | | | smtO-Bxb1 | JC605 | 0.6 |

Supplementary Table 8: Concentration of aTFs proteins for cell-free reaction in Figure 3C and 3D

| aTFs Protein | aTFs Protein Number | Recombinase | aTFs Protein Conc. (uM) | Chemical | Chemical Conc. (nM) |
|---|---|---|---|---|---|
| tetR | JC20 | tetO-Cre | 12.5 | Tetracycline | 12.5 |
| | | tetO-PhiC | 1.25 | Tetracycline | 2.5 |
| | | tetO-Bxb1 | 1.25 | Tetracycline | 2.5 |
| smtB | JC22 | smtO-Cre | 10 | Zinc | 12.5 |
| | | smtO-PhiC | 10 | Zinc | 50 |
| | | smtO-Bxb1 | 10 | Zinc | 125 |

Supplementary Table 9: Concentration of DNA for cell-free reaction in Figure 4B

| Reporter | Plasmid Number | Plasmid Conc. (nM) | Recombinase | Plasmid Number | Plasmid Conc. (nM) |
|---|---|---|---|---|---|
| NOR | JC305 | 6 | tetO-Cre | JC287 | 0.6 |
| AND | JC306 | 6 | smtO-PhiC | JC604 | 0.6 |
| NAND | JC431 | 6 | | | |
| A IMPLY B | JC434 | 6 | | | |
| NOT A | JC432 | 6 | | | |
| NOT B | JC433 | 6 | | | |
| A NIMPLY B | JC436 | 6 | | | |
| B NIMPLY A | JC437 | 6 | | | |

Supplementary Table 10: Concentration of aTFs proteins for cell-free reaction in Figure 4B and 5B

| aTFs Protein | aTFs Protein Number | aTFs Protein Conc. (uM) | Chemical | Chemical Conc. (nM) |
|---|---|---|---|---|
| tetR | JC20 | 12.5 | Tetracycline | 12.5 |
| smtB | JC22 | 5 | Zinc | 12.5 |

Supplementary Table 11: Concentration of DNA for cell-free reaction in Figure 5B

| Reporter | Plasmid Number | Plasmid Conc. (nM) | Recombinase | Plasmid Number | Plasmid Conc. (nM) |
|---|---|---|---|---|---|
| 2 Input Decoder | JC493 | 6 | tetO-Cre | JC287 | 0.6 |
| | | | smtO-PhiC | JC604 | 0.6 |

Supplementary Table 12: Concentration of DNA for cell-free reaction in Figure 6B

| Reporter | Plasmid Number | Plasmid Conc. (nM) | Recombinase | Plasmid Number | Plasmid Conc. (nM) |
|---|---|---|---|---|---|
| Cre | JC213 | 6 | Cre | JC40 | 0.6 |

Supplementary Table 13: Concentration of DNA for cell-free reaction in Supplementary Figure 1A

| Reporter | Plasmid Number | Plasmid Conc. (nM) | Recombinase | Plasmid Number | Plasmid Conc. (nM) |
|---|---|---|---|---|---|
| Reporter | JC42 | 6 | Cre | JC40 | 0.6 |
| LoxP-mCherry | JC466 | 6 | | | |

Supplementary Table 14: Concentration of DNA for cell-free reaction in Supplementary Figure 1B

| Reporter | Plasmid Number | Plasmid Conc. (nM) | Recombinase | Plasmid Number | Plasmid Conc. (nM) |
|---|---|---|---|---|---|
| Cre | JC213 | 6 | Cre | JC40 | 0.6 |
| PhiC | JC196 | 6 | PhiC | JC138 | 0.6 |
| Bxb1 | JC271 | 6 | Bxb1 | JC140 | 0.6 |
| A118 | JC311 | 6 | A118 | JC348 | 0.6 |
| TP901 | JC312 | 6 | TP901 | JC350 | 0.6 |
| VCre | JC315 | 6 | VCre | JC352 | 0.6 |

Supplementary Table 15: Concentration of DNA for cell-free reaction in Supplementary Figure 1C

| Reporter | Plasmid Number | Plasmid Conc. (nM) | Recombinase | Plasmid Number | Plasmid Conc. (nM) |
|---|---|---|---|---|---|
| Cre | JC213 | 6 | Cre | JC40 | 0-6 |
| PhiC | JC196 | 6 | PhiC | JC138 | 0-6 |
| Bxb1 | JC271 | 6 | Bxb1 | JC140 | 0-6 |

Supplementary Table 16: Concentration of DNA for cell-free reaction in Supplementary Figure 2

| Reporter | Plasmid Number | Plasmid Conc. (nM) | Recombinase | Plasmid Number | Plasmid Conc. (nM) |
|---|---|---|---|---|---|
| Original | JC42 | 6 | Cre | JC40 | 0.6 |
| Terminator Characterized | JC465 | 6 | | | |
| mutLoxP | JC757 | 6 | | | |
| Insert Length Characterized | JC213 | 6 | | | |

Supplementary Table 17: Concentration of DNA for cell-free reaction in Supplementary Figure 3A

| Reporter | Plasmid Number | Plasmid Conc. (nM) | Recombinase | Plasmid Number | Plasmid Conc. (nM) |
|---|---|---|---|---|---|
| Cre | JC213 | 6 | tetO-Cre | JC287 | 0.6 |
| PhiC | JC196 | 6 | tetO-PhiC | JC288 | 0.6 |
| Bxb1 | JC271 | 6 | tetO-Bxb1 | JC289 | 0.6 |

Supplementary Table 18: Concentration of aTFs proteins for cell-free reaction in Figure 3A

| aTFs Protein | v Protein Number | Recombinase | aTFs Protein Conc. (uM) | Chemical | Chemical Conc. (nM) |
|---|---|---|---|---|---|
| tetR | JC20 | tetO-Cre | 0.125, 1.25, 12.5 | Tetracycline | 0-62.5 |
| | | tetO-PhiC | 0.125, 1.25, 12.5 | Tetracycline | 0-62.5 |
| | | tetO-Bxb1 | 0.125, 1.25, 12.5 | Tetracycline | 0-62.5 |

Supplementary Table 19: Concentration of DNA for cell-free reaction in Supplementary Figure 4A

| Reporter | Plasmid Number | Plasmid Conc. (nM) |
|---|---|---|
| HucO-mCherry | JC303 | 6 |
| QacA-mCherry | JC301 | 6 |
| OrtO-mCherry | JC297 | 6 |
| CtcO-mCherry | JC298 | 6 |
| TtgO-mCherry | JC302 | 6 |

Supplementary Table 20: Concentration of proteins and chemicals for cell-free reaction in Supplementary Figure 4A

| aTFs Protein | aTFs Protein Number | aTFs Protein Conc. (uM) | Chemicals | Chemicals Conc. (nM) |
|---|---|---|---|---|
| HucR | JC309 | 68.8 | Uric Acid | 500 |
| QacR | JC307 | 2.25 | Benzalkonium Chloride | 100 |
| OtrR | JC316 | 25 | Oxytetracycline | 50 |
| CtcS | JC317 | 12.5 | Chlortetracycline | 12.5 |
| TtgR | JC308 | 75 | Naringenin | 250 |

Supplementary Table 21: Concentration of DNA for cell-free reaction in Supplementary Figure 4B

| Reporter | Plasmid Number | Plasmid Conc. (nM) | Recombinase | Plasmid Number | Plasmid Conc. (nM) |
|---|---|---|---|---|---|
| Cre | JC213 | 6 | HucO-Cre | JC338 | 0.6 |
| PhiC | JC196 | 6 | HucO-PhiC | JC339 | 0.6 |
| Bxb1 | JC271 | 6 | HucO-Bxb1 | JC340 | 0.6 |

Supplementary Table 22: Concentration of protein and chemicals for cell-free reaction in Supplementary Figure 4B

| aTFs Protein | aTFs Protein Number | Recombinase | aTFs Protein Conc. (uM) | Chemical | Chemical Conc. (nM) |
|---|---|---|---|---|---|
| HucR | JC309 | HucO-Cre | 68.8 | Uric Acid | 500 |
| | | HucO-PhiC | 68.8 | Uric Acid | 500 |
| | | HucO-Bxb1 | 68.8 | Uric Acid | 500 |

Supplementary Table 23: Concentration of DNA for cell-free reaction in Supplementary Figure 5 and Supplementary Figure 6

| Reporter | Plasmid Number | Plasmid Conc. (nM) | Recombinase | Plasmid Number | Plasmid Conc. (nM) |
|---|---|---|---|---|---|
| NOR | JC305 | 6 | Cre | JC40 | 0.6 |
| AND | JC306 | 6 | PhiC | JC138 | 0.6 |
| NAND | JC431 | 6 | | | |
| A IMPLY B | JC434 | 6 | | | |
| NOT A | JC432 | 6 | | | |
| NOT B | JC433 | 6 | | | |
| A NIMPLY B | JC436 | 6 | | | |
| B NIMPLY A | JC437 | 6 | | | |

Supplementary Table 24: Concentration of DNA for cell-free reaction in Supplementary Figure 7

| Reporter | Plasmid Number | Plasmid Conc. (nM) | Recombinase | Plasmid Number | Plasmid Conc. (nM) |
|---|---|---|---|---|---|
| 2 Input Decoder | JC493 | 6 | Cre | JC40 | 0.6 |
| | | | PhiC | JC138 | 0.6 |
